# What is Slough? A pilot study to define the proteomic and microbial composition of wound slough and its implications for wound healing

**DOI:** 10.1101/2023.11.02.565225

**Authors:** Elizabeth C. Townsend, J. Z. Alex Cheong, Michael Radzietza, Blaine Fritz, Matthew Malone, Thomas Bjarnsholt, Karen Ousey, Terry Swanson, Gregory Schultz, Angela L.F. Gibson, Lindsay R. Kalan

**Affiliations:** Department of Medical Microbiology and Immunology, University of Wisconsin School of Medicine and Public Health, Madison, WI, United States; Microbiology Doctoral Training Program, University of Wisconsin-Madison, Madison, WI, United States; Medical Scientist Training Program, University of Wisconsin School of Medicine and Public Health, Madison, WI, United States; Infectious Diseases and Microbiology, Western Sydney University, Australia; Department of Immunology and Microbiology, University of Copenhagen, Denmark; Department of Clinical Microbiology, Copenhagen University Hospital, Rigshospitalet, Denmark; International Wound Infection Institute, London, United Kingdom; Institute of Skin Integrity and Infection Prevention, University of Huddersfield, United Kingdom; Obstetrics and Gynecology, University of Florida, United States; Department of Surgery, University of Wisconsin School of Medicine and Public Health, Madison, WI, United States; M.G. DeGroote Institute for Infectious Disease Research, David Braley Centre for Antibiotic Discovery, Department of Biochemistry and Biomedical Sciences, McMaster University, Hamilton, Ontario, Canada

## Abstract

Slough is a well-known feature of non-healing wounds. This study aims to determine the proteomic and microbiologic components of slough as well as interrogate the associations between wound slough components and wound healing. Twenty-three subjects with slow-to-heal wounds and visible slough were enrolled. Etiologies included venous stasis ulcers, post-surgical site infections, and pressure ulcers. Patient co-morbidities and wound healing outcome at 3-months post-sample collection was recorded. Debrided slough was analyzed microscopically, through untargeted proteomics, and high-throughput bacterial 16S-ribosomal gene sequencing. Microscopic imaging revealed wound slough to be amorphous in structure and highly variable. 16S-profiling found slough microbial communities to associate with wound etiology and location on the body. Across all subjects, slough largely consisted of proteins involved in skin structure and formation, blood-clot formation, and immune processes. To predict variables associated with wound healing, protein, microbial, and clinical datasets were integrated into a supervised discriminant analysis. This analysis revealed that healing wounds were enriched for proteins involved in skin barrier development and negative regulation of immune responses. While wounds that deteriorated over time started off with a higher baseline Bates-Jensen Wound Assessment Score and were enriched for anerobic bacterial taxa and chronic inflammatory proteins. To our knowledge, this is the first study to integrate clinical, microbiome, and proteomic data to systematically characterize wound slough and integrate it into a single assessment to predict wound healing outcome. Collectively, our findings underscore how slough components can help identify wounds at risk of continued impaired healing and serves as an underutilized biomarker.

## Introduction

Chronic, non-healing wounds impose a significant, underappreciated burden to affected individuals and the healthcare system. An estimated 2 – 10% of the general population in Australia, the United Kingdom and the United States suffer from chronic wounds.^1–4^ Individuals with conditions known to impair wound healing, such as peripheral arterial disease, venous insufficiency, immune-compromised, obesity, diabetes, impaired sensation, and spinal cord injuries are at the highest risk for developing chronic wounds.^5^ With the prevalence of these comorbidities on the rise, chronic wounds are anticipated to pose a growing burden for patients and the healthcare system.^2,3^ Thus, identifying biomarkers to distinguish chronic wounds that are likely to heal from those that may benefit from intensive therapies to promote healing is a critical imperative.

A hallmark feature of chronic wounds is the presence of slough, which mainly consists of devitalized tissue that overlays the wound bed. Slough is hypothesized to arise as a byproduct of prolonged wound inflammation.^6–8^ On a macro level, slough has highly variable physical characteristics ranging in consistency, color, odor, and attachment to the wound bed even across a single wound’s surface.^6,8^ Subsequently the appearance varies widely wound-to-wound and patient-to-patient. Although, to date, there are no studies interrogating slough directly, assessments of exudative fluid from surface of chronic wounds and wound biopsies suggest that the wound surface and associated slough is enriched for various types of collagen, extracellular matrix proteins, matrix metalloproteases, and proteins related to inflammatory immune responses.^9–14^ Slough can also be infiltrated by an array of bacterial either as single cells or by forming aggregates and biofilm.^15–20^ However, due to the highly variable appearance and inconsistencies in even defining slough between providers, it is difficult to distinguish slough with or without microbial biofilm from infected wound exudate.^15,21^ Ultimately, slough’s variable nature has led to inconsistent clinical approaches to wound management.

One dominant theory proposes that slough inhibits wound healing by prolonging the inflammatory phase of healing, preventing the formation of granulation tissue and subsequent wound contraction. Slough is commonly associated with biofilm, although limited evidence exists to support the idea that slough is primarily microbial in nature. Slough may serve as a reservoir attractive to bacteria on the wound bed that subsequently promotes biofilm formation, however this is also challenging to quantify.^6^ In the absence of conclusive data, standard chronic wound care focuses on proper debridement to remove devitalized tissue, reduce potential surface microbial burden, and ideally return the wound to an acute state to stimulate tissue repair.^22^ However, less than 50% of wounds respond or go on to heal following debridement.^23^ Conversely, some wounds with slough present will heal without debridement, suggesting that the presence of slough does not always indicate that healing is disrupted.^8^

Despite it being a common feature of chronic wounds, a detailed molecular characterization of the host and microbial components within slough from different wound etiologies is missing. A systematic analysis of slough composition and factors associated with wound healing outcomes could shape wound treatment strategies and aide in triaging high risk patients into specialty care.

With this pilot study, we aim to characterize the human and microbial components of slough collected from wounds of various etiologies. Collectively we show that wound slough is primarily composed of proteins associated with the structure and formation of the skin, blood clot formation, and various immune responses. Wound slough is highly polymicrobial and exhibits signatures associated with both wound etiology and location on the body. Finally, slough protein profiles from wounds with a healing trajectory are significantly different than slough protein profiles from non-healing wounds, suggesting they may serve as a prognostic marker. Rather than being discarded, slough may be a critical indicator to predict if a wound is more likely on the trajectory toward healing or at risk of deteriorating.

## Methods

### Subject Identification and Enrollment

Adults 18 years or older with chronic wounds were recruited from UW-Health Wound Care Clinics under an IRB approved protocol (Study ID: 2020-1002). Examples of wounds identified for possible inclusion included and were not limited to, chronic or non-healing diabetic ulcers, pressure ulcers, venous ulcers, surgical or procedural wounds, trauma wounds, burn wounds, and wounds of unknown or other etiology. On the day of sample collection, subject wounds were measured, evaluated and scored according to the Bates-Jensen Wound Assessment Tool.^24^ Information related to the wound’s etiology and care, wound measurements from the most recent previous visit, and patient co-morbidities were extracted from the medical record. Digital photos of the wound were taken before and after the debridement procedure. Swabs for microbiome analysis of the wound edge and center were collected using Levine’s technique and placed into 300 μl of DNA/RNA Shield (Zymo Research, Irvine, CA) and stored at –80°C until further processing. Swabs were spun down using DNA IQ Spin Baskets (Promega, Madison, WI) and DNA was extracted. Swabs designated for microbial culture were taken from the wound center using Levine’s technique into 1 ml of liquid Amies (Copan Diagnostics Inc., Murrieta, CA). Swabs were stored at 4°C for less than 2 hours before being processed for microbial culture.

All subjects received sharp debridement of their wounds. Prior to debridement wounds were washed with soap and rinsed with water. Debridement was performed by a skilled practitioner with surgical instruments such as scalpel, curette, scissors, rongeur, and/ or forceps. Removed slough material was collected into 1ml of DNA/RNA Shield (Zymo Research, Irvine, CA) and stored at 4°C before sectioning for scanning electron microscopy (SEM), fluorescence in situ hybridization (FISH), and proteomics. Remaining slough material was stored at –80°C.

Samples from South Western Sydney Hospital were collected and processed as described by Malone et al.^25^ Adults 18 years or older presenting with a diabetes-related foot ulcer with visible signs of slough were recruited for the study. The collection of samples and their corresponding patients was undertaken as a sub-analysis of a larger clinical study, with samples being obtained following written consent. Ethics approval for the larger clinical study and the slough sub-analysis was approved by South Western Sydney LHD Research and Ethics Committee. All DFUs were debrided and rinsed with 0.9% NaCl prior to specimen collection. For DNA sequencing, patient wound slough was removed from the ulcer base with a dermal curette and immediately stored in RNA Shield (Zymo Research, Irvine, CA) at 4°C for 24 hours before being frozen at –80°C until further processing. For PNA-FISH, tissue specimens were obtained through a dermal ring curette from the wound bed of each DFU. Following removal, tissue specimens were rinsed vigorously in a phosphate buffer solution (PBS) bath to remove any coagulated blood and to reduce the number of planktonic microorganisms. Tissue specimens were immediately fixed in 4% paraformaldehyde overnight at 4°C, then transferred into 70% ethanol and stored at –20°C

### Microbial Culture and Bacterial Isolate Identification

Swabs designated for microbial culture were spun down using DNA IQ Spin Baskets (Promega, Madison, WI). A portion of each sample was serially diluted with 1X phosphate buffered saline and plated onto Tryptic Soy Agar (TSA) with 5% sheep blood (BBL, Sparks, MD) for quantitative bacterial culture. Plates were incubated at 35°C overnight. To isolate culturable bacteria, colonies with distinct morphology were isolated and incubated at 35°C overnight on TSA with 5% sheep blood then single colonies were inoculated into liquid Tryptic Soy Broth (TSB) for overnight incubation. To identify each bacterial isolate, a portion of the overnight TSB culture underwent DNA extraction and sanger sequencing (Functional Biosciences, Madison, WI) of the bacterial 16S ribosomal RNA gene. The remaining portion of the isolate culture was stored in glycerol at –80°C.

### DNA/RNA extraction, library construction, sequencing

DNA extraction on samples collected in the USA was performed as previously described with minor modifications.^26^ Briefly, 300 μl of yeast cell lysis solution (from Epicentre MasterPure Yeast DNA Purification kit), 0.3 μl of 31,500 U/μl ReadyLyse Lysozyme solution (Epicentre, Lucigen, Middleton, WI), 5 μl of 1 mg/ml mutanolysin (M9901, Sigma-Aldrich, St. Louis, MO), and 1.5 μl of 5 mg/ml lysostaphin (L7386, Sigma-Aldrich, St. Louis, MO) was added to 150 μl of swab liquid before incubation for one hour at 37°C with shaking. Samples were transferred to a tube with 0.5 mm glass beads (Qiagen, Germantown, Maryland) and bead beat for 10 min at maximum speed followed by a 30 min incubation at 65°C with shaking, 5 min incubation on ice. The sample was spun down at 10,000 rcf for 1 min and the supernatant was added to 150 μl of protein precipitation reagent (Epicentre, Lucigen, Middleton, WI). Remaining steps followed the recommended PureLink Genomic DNA Mini Kit (Invitrogen, Waltham, MA) protocol for DNA extraction and purification. 16S rRNA gene amplicon libraries targeting the V1-V3 or V4 region were constructed using a dual-indexing method and sequenced on a MiSeq with a 2×300 bp run format (Illumina, San Diego, CA). Reagent-only negative controls were carried through the DNA extraction and sequencing process.

Swabs obtained from DFUs in Australia were defrosted on ice prior to DNA extraction. Genomic DNA was extracted using Qiagen DNeasy PowerBiofilm kit (Cat No./ID: 24000-50) following the manufacturer’s instructions. Preparation of the16S library and DNA sequencing was carried out by a commercial laboratory (Ramaciotti Centre for Genomics, University of New South Wales, Australia) on the Illumina MiSeq platform (2×300bp) targeting the V1-V3 (27f/519r) 16S region.

### Sequence analysis

The QIIME2^27^ environment was used to process DNA-based 16S rRNA gene amplicon data. Paired end reads were trimmed, quality filtered, and merged into amplicon sequence variants (ASVs) using DADA2. Taxonomy was assigned to ASVs using a naive Bayes classifier pre-trained on full length 16S rRNA gene 99% operational taxonomic unit (OTU) reference sequences from the Greengenes database (version 13_8). Using the qiime2R package, data was imported into RStudio (version 1.4.1106) running R (version 4.1.0) for further analysis using the phyloseq package.^28^ Negative DNA extraction and sequencing controls were evaluated based on absolute read count and ASV distribution in true patient samples. Abundances were normalized proportionally to total reads per sample. Data was imported into RStudio running R (version 4.2.1) for analysis. Relative abundance plots were produced using the package ggplot2, where taxa below 1% relative abundance were pooled into an “Other” category.

### Proteomics

Debrided slough tissue samples were weighed and placed in PowerBead tubes containing 1.4mm ceramic beads (Qiagen, Germantown, Maryland) for tissue homogenization, proteomic processing, and analysis at the University of Wisconsin Mass Spectrometry and Proteomics Core Facility. In brief, samples were labeled and pooled for multiplex relative mass spectrometry (MS) quantification with the TMTpro 16plex labeling kit (ThermoFisher Scientific, Waltham, MA) and underwent Liquid Chromatography with tandem mass spectrometry on an Orbitrap Elite mass spectrometer (ThermoFisher Scientific). Protein sequences were matched to known human and bacterial proteins. Functions associated with each protein were gathered from the Gene Ontology (GO) database, KEGG Pathways, Reactome Pathways, and WikiPathways databases. Data was imported into RStudio for analysis. To determine the most enriched proteins and their associated biologic processes within slough, abundances were normalized proportionally to total abundance per sample and the ranked dataset was analyzed via the Gene Ontology enRIchment analysis (GORILA) and visualization tool.^29^ Differential protein expression between subject groups was assessed via DEqMS.^30^ To determine the key biologic processes for smaller sets of proteins, such as those enriched within subject groups or the proteins within each of the k-means protein clusters (see *Integration of Biologic Data Sets* below), small sets of proteins were submitted as unranked lists to the GO Enrichment Analysis tool.^31,32^

### Fluorescence in situ hybridization (FISH)

Formalin-fixed paraffin-embedded (FFPE) histological sections were deparaffinized in xylene and rehydrated in a series of ethanol washes (100%, 99%, 95%, and 0%). Subsequently, the samples were allowed to hybridize at 46°C for 4 hours in hybridization solution (900 mM NaCl, 20 mM Tris pH 7.5, 0.01% SDS, 20% formamide, 2 μM FISH probe). The FISH probe used was a DNA oligonucleotide (EUB388 sequence) with a 3’-conjugated TEX615 fluorophore (Integrated DNA Technologies, Coralville, IA, USA). Samples were washed in excess wash buffer (215 mM NaCl, 20 mM Tris pH 7.5, 5 mM EDTA) at 48°C for 15 mins, dipped into ice cold water, 100% ethanol, drained, and air-dried. Slides were mounted with Prolong Glass antifade mounting medium with NucBlue counterstain (Thermo Fisher Scientific, Waltham, MA, USA) and a glass coverslip of #1.5 thickness and stored flat to cure overnight in the dark. Micrographs were acquired using a Zeiss 780 confocal laser scanning microscope on the red TEX615, blue Hoescht, and green GFP (tissue autofluorescence) channels using 5× and 63× objectives. Zeiss Zen software was used to analyze tiled images, z-stacks, and generate maximum intensity projections.

### PNA-FISH

As described by Nadler et al.^33^, formalin-fixed, paraffin-embedded samples were cut, deparaffinized and rehydrated following standard procedures. Subsequently, the samples were stained with a PNA-FISH-TexasRed-5-conjugated universal bacterial (BacUni) 16s rRNA probe (AdvanDx, Woburn, MA, US), incubated and then counterstained with 3 µM 4′,6-diamidino-2-phenylindole (DAPI) (life Technologies, Eugene, OR, USA). The samples were afterward mounted (ProLong™ Gold Antifade Mountant, Life Technologies) and a coverslip was added (Marienfield, Lauda-Königshoffen, Germany). Slides were evaluated using a CLSM (Axio Imager.Z2, LSM880 CLSM; Zeiss, Jena, Germany). Images were taken using 405 nm (DAPI) and 561 nm (TexasRed-5) lasers, as well as a 488 nm laser for visualizing the green autofluorescence of the surrounding tissue. Images were subsequently processed with IMARIS 9.2 (Bitplane, Zurich, Switzerland) and presented using “Easy 3D”.

### Scanning Electron Microscopy

Wound slough specimens were rinsed with PBS and fixed overnight in 5 mL of 1.5% glutaraldehyde in 0.1 M sodium phosphate buffer (pH 7.2) at 4°C. Samples were rinsed, treated with 1% osmium tetroxide for 1 h, and then washed again in buffer. Samples were dehydrated through a series of ethanol washes (30–100%) followed by critical point drying (14 exchanges on low speed) and were subsequently mounted on aluminum stubs with a carbon adhesive tab and carbon paint. Samples were left to dry in a desiccator overnight. Following sputter coating with platinum to a thickness of 20 nm, samples were imaged in a scanning electron microscope (Zeiss LEO 1530-VP) at 3 kV.

### Integration of Biologic Data Sets

To reduce the complexity of the proteomics data for integrative analysis protein abundances were normalized, mean centered and grouped via k-means clustering. The optimal number of protein clusters was determined by the Gap-Statistics method. Since there was no significant difference in the microbial community composition at the wound edge or center, taxa relative abundances from the wound edge and center were averaged to create a summative wound slough microbiome for each subject. The 15 microbial ASVs with greater than 1% relative abundance in at least two summative subject slough microbiome samples were included for this analysis. To predict the variables associated with wound healing, the protein cluster, summative slough microbiome, and the numerical Bates-Jensen Wound Assessment score datasets were integrated into a supervised Partial Least Squares – Discriminant Analysis (PLS-DA, aka. Data Integration Analysis for Biomarker discovery using Latent variable approaches for Omics studies [DIABLO]) via the via MixOmics^34^ R-studio package.

### Statistical analyses

Statistical analyses were conducted in R studio running R (version 4.2.1). Bates Jensen wound assessment scores were analyzed via Prism (version 9.2.0).

### Data availability

Sequence reads for this project can be found under NCBI BioProject PRJNA1021648. Code for analysis and generation of figures can be found on GitHub at https://github.com/Kalan-Lab/Townsend_etal_WhatIsSlough.

## Results

### Subject and Wound Characteristics

Twenty-three subjects with wounds of various etiologies were included in this study. To address potential inconsistencies between sites and batch effects, the main analysis focuses on ten patients recruited in the United States (Table 1, Table S1). Data from the remaining thirteen patients is available in supplementary materials. For the ten patients, prior to sample collection wounds were measured, evaluated and scored according to the Bates-Jensen Wound Assessment Tool (Table 2, Table S1).^35^ Overall Bates-Jensen Wound Assessment scores ranged from 26 to 46 (mean 37.4) out of 60 points with higher scores indicating greater wound degeneration. Photos of the wounds before sharp debridement of the superficial wound slough are in supplemental figure 1.

**Table 1:**
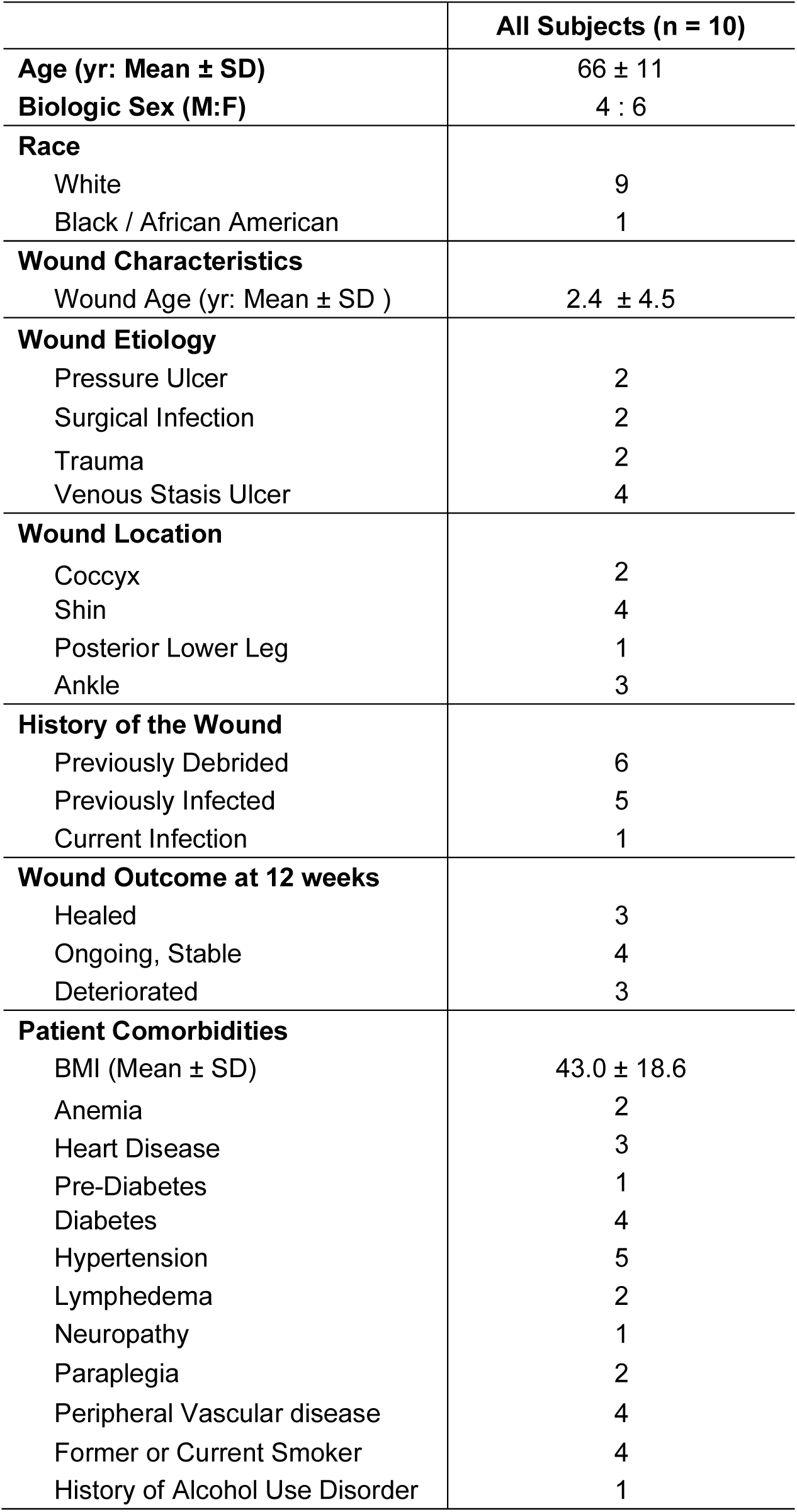
Subject and Wound Characteristics.

**Table 2:** Bates-Jensen Wound Assessment Scores By Wound Healing Outcome at 3 months. Data represented as mean +/− standard deviation

|  | **Healed**<br>**(n=3)** | **Ongoing**<br>**(n=4)** | **Deteriorated (n =**<br>**3)** | **All Subjects (n =**<br>**10)** |
| --- | --- | --- | --- | --- |
| **Size** | 2.00 ± 1.00 | 3.25 ± 1.71 | 3.00 ± 2.00 | 2.80 ± 1.55 |
| **Depth** | 2.67 ± 0.58 | 2.75 ± 0.5 | 4.00 ± 1.00 | 3.10 ± 0.88 |
| **Edges** | 2.67 ± 1.15 | 3.25 ± 0.5 | 3.67 ± 0.58 | 3.20 ± 0.79 |
| **Undermining** | 2.33 ± 2.31 | 1.50 ± 1.00 | 2.33 ± 2.31 | 2.00 ± 1.70 |
| **Necrotic Tissue Type** | 2.67 ± 0.58 | 3.00 ± 0.82 | 3.00 ± 1.00 | 2.90 ± 0.73 |
| **Necrotic Tissue Amount** | 2.67 ± 0.58 | 2.75 ± 0.5 | 2.67 ± 2.08 | 2.70 ± 1.06 |
| **Exudate Type** | 2.67 ± 0.58 | 3.25 ± 0.5 | 3.67 ± 1.15 | 3.20 ± 0.79 |
| **Exudate Amount** | 3.33 ± 0.58 | 3.50 ± 0.58 | 4.33 ± 0.58 | 3.70 ± 1.32 |
| **Skin Color Surrounding**<br>**Wound** | 1.00 ± 0 | 1.75 ± 1.50 | 2.67 ± 1.53 | 1.80 ± 1.32 |
| **Edema** | 1.67 ± 1.15 | 2.50 ± 1.73 | 2.33 ± 1.53 | 2.20 ± 1.40 |
| **Induration** | 1.00 ± 0 | 1.75 ± 0.96 | 2.00 ± 1.73 | 1.60 ± 1.08 |
| **Granulation** | 2.00 ± 1.00 | 3.75 ± 0.50 | 4.00 ± 0 | 3.30 ± 1.06 |
| **Epithelialization** | 5.00 ± 0 | 4.75 ± 0.50 | 5.00 ± 0 | 4.90 ± 0.32 |
| **Total Score** | 31.67 ± 4.93 | 37.75 ± 3.86 | 42.67 ± 3.06 | 37.4 ± 5.7 |

Wound status at 3 months post-sample collection was recorded (Table 1). At this time, 3 of the subjects’ wounds healed, 4 were ongoing yet stable in size and clinical assessment, and 3 wounds had deteriorated (e.g. significantly increased in size, depth, and/or continued antibiotic resistant infection). The total Bates-Jensen Wound Assessment Score, and several of the sub-scores trended higher in wounds that deteriorated compared to those that went on to heal (p-values < 0.1, yet > 0.05, Mann-Whitney t-test. Table 2). However, none of these comparisons reached statistical significance, likely due to the relatively small number of subjects within each group.

### Slough protein composition is associated with wound age and healing trajectory

Slough tissue was first characterized by untargeted proteomics to determine the overall protein composition (Table S2). 11,058 peptide fragments (7,302 unique peptide groups) corresponding to 1,447 unique protein signatures were detected. To identify the biologic processes, molecular functions, and cellular components that associated with protein features, abundant proteins identified across all samples were analyzed using the Gene Ontology enRIchment analysis (GORILA) and visualization tool.^29^ This demonstrated that wound slough is enriched for proteins derived from both intracellular and extracellular components, and notably enriched for proteins specific to skin tissue, such as the cornified envelope and keratin filaments (Fig. 1, Fig. S2, Table S3). Molecular pathway analysis determined slough samples are enriched for proteins involved in ion and metabolite binding. This analysis further confirmed that wound slough is significantly enriched for proteins involved in skin barrier formation, wound healing, blood clotting, as well as various immune functions including responding to bacteria, acute inflammatory responses, immune effector cell responses, and humoral immunity (Fig. 1).

**Figure 1:**
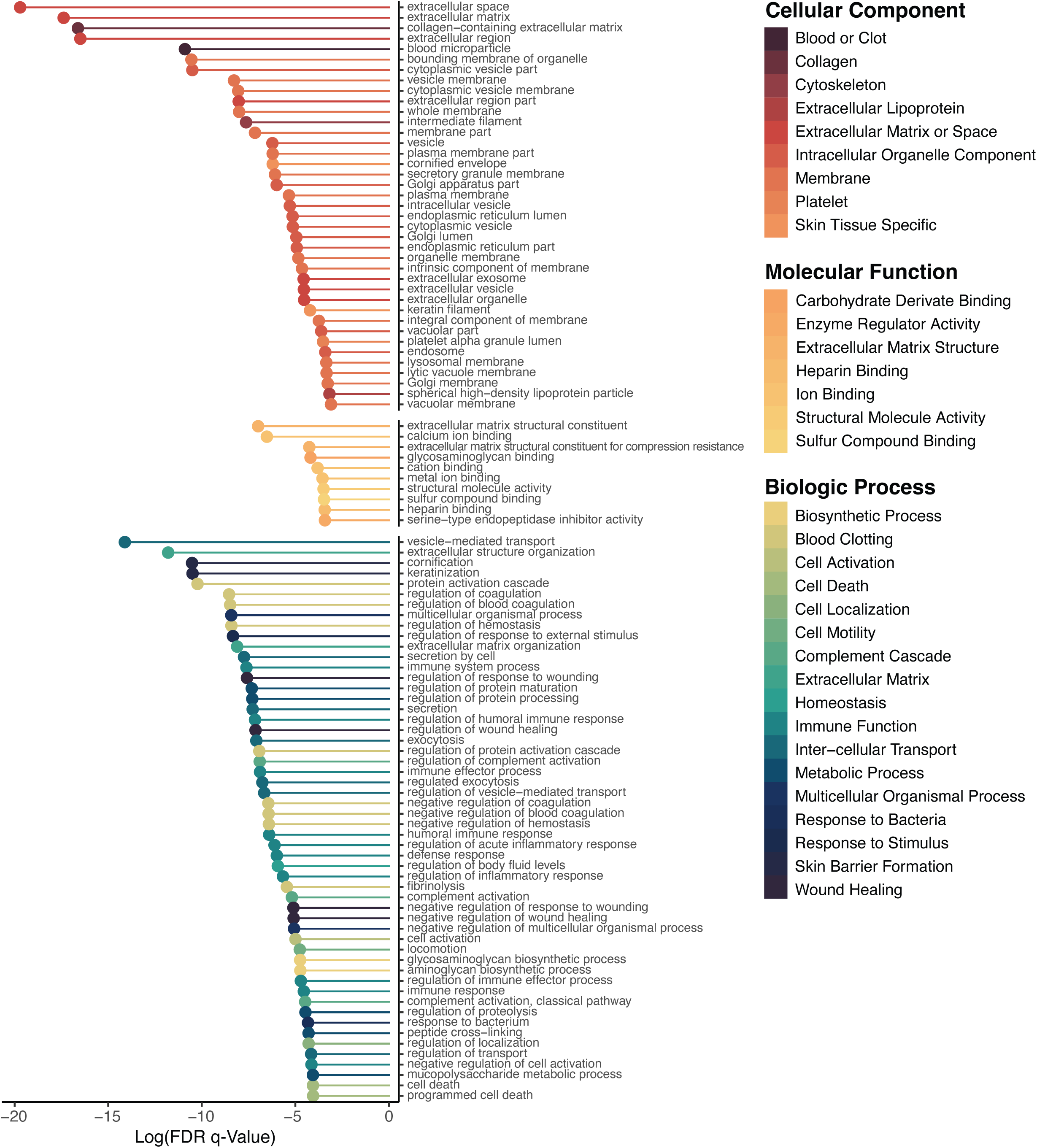
Wound slough is enriched for proteins involved in skin barrier formation, wound healing, blood clotting, and various immune functions including responding to bacteria. Debrided slough tissue was sent for proteomic characterization via mass spectrometry. The most abundant proteins across all samples were input as a ranked list to the Gene Ontology enRIchment analysis (GORILA) and visualization tool.^29^ Significantly enriched GO terms are listed by their description and ordered by their FDR-qValue. GO terms associated with extracellular and cellular components are in the top section (reds), those associated with molecular functions are in the middle (oranges to yellows), and those associated with biologic processes are in the bottom section (yellow-greens to blues). Associated color-coded trimmed directed acyclic graphs (DAG) of all significantly enriched GO terms as grouped by component, function, and biologic process are in supplemental figure 2. More detail, including GO term annotations, descriptions, enrichment, number of proteins (Uniprot Genes) involved from our dataset involved in each GO Term, and FDR-qValues are in supplemental table 3.

To determine if the protein composition of slough is associated with clinical features and wound healing outcomes, hierarchical clustering using Euclidean distances was performed to identify patterns across the dataset. Clustering appeared to be driven by the wound age at sample collection and clinical outcome at 12 weeks-post collection defined as healed, ongoing but stable, or deteriorating (Fig. 2A). Proteins differentially abundant in healing wounds compared to those that were stable or deteriorated, were then determined using DEqMS.^30^ Forty-eight proteins were differently abundant between healing wounds and deteriorating wounds, while thirty-two proteins were differently abundant between healing wounds and those that were ongoing yet stable (Fig. 2B-D, Table S4). GO Enrichment Analysis^32^ shows that healing wounds are enriched for proteins involved in skin barrier development (e.g. cornifin-B, and 14-3-3 protein sigma), wound healing (e.g. beta-2-glycoprotein 1), blood clot formation (e.g. coagulation factor XIII) and responses to bacteria and external stress (e.g. immunoglobins, cystatin-F, and peroxiredoxin-6). Conversely, deteriorating wounds are enriched for proteins involved in immune responses categorized as chronic inflammatory responses (e.g. AP-1 complex subunit gamma-1 and NLR family proteins) and the compliment cascade (e.g. Complement factor H). Finally, differential protein analysis between newer or older wounds found newer wounds (defined as being present for less than 1 year) are enriched in proteins involved in epithelial barrier formation and integrity (e.g. epithelial cell division and epithelial cell-to-cell adhesion), neutrophil degranulation, and response to bacteria (Fig. S3).

**Figure 2:**
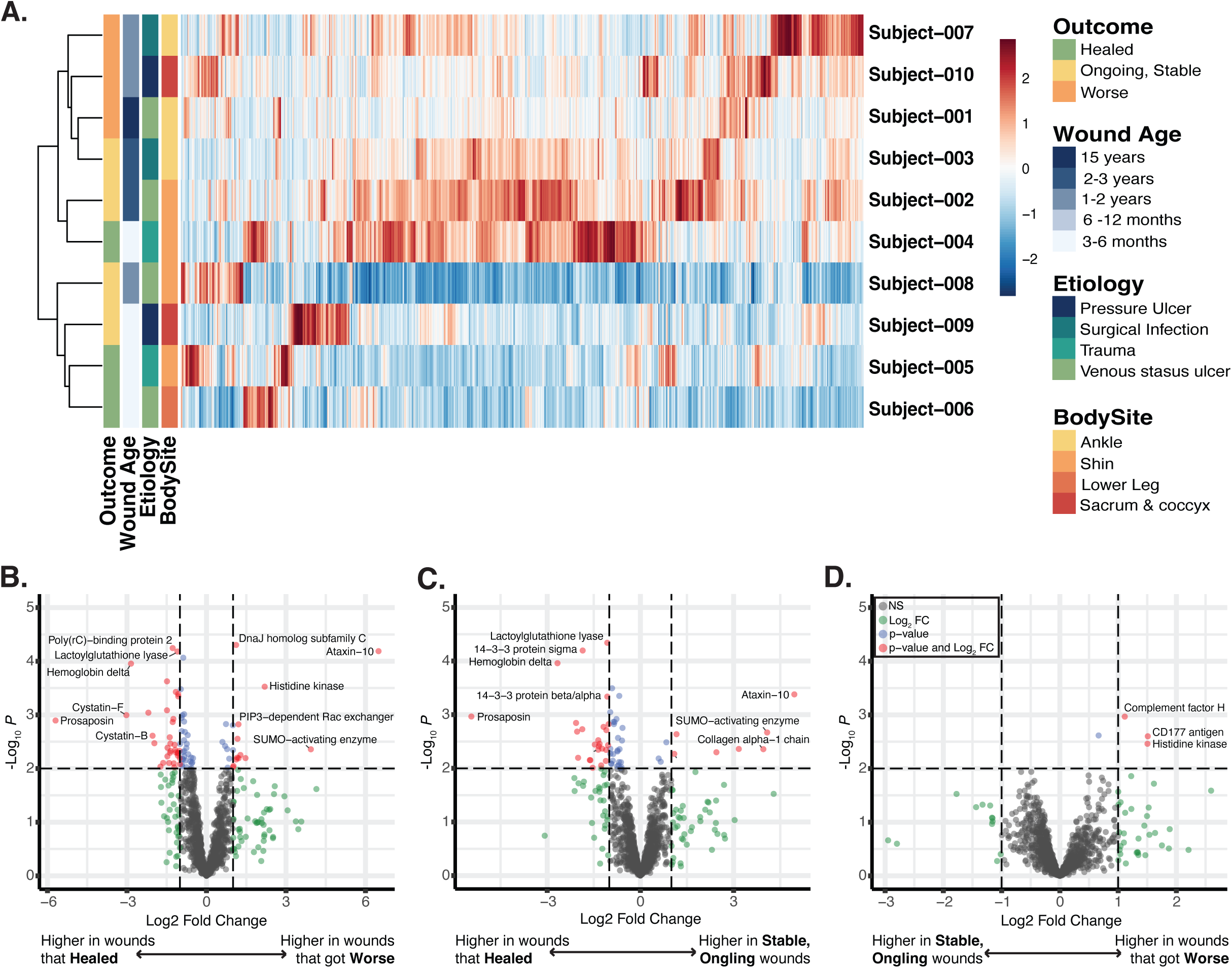
Wounds that go on to heal, are ongoing, or deteriorate are enriched for different proteins. A) heat map demonstrating each proteins’ relative expression across all subjects demonstrates that samples largely grouped by the wounds relative age and outcome 3 months after the sample collection (e.g whether the wound went on to heal, is ongoing but stable, or continued to deteriorate). B-D) Subjects were grouped by the wound’s outcome and groups were assessed for differential protein expression via DEqMS. Volcano plots indicating the proteins with significantly greater expression in; B) wounds that continued to deteriorate vs. wounds that went on to heal; C) wounds that were stable but on going vs. those that healed: D) and wounds that continued to deteriorate vs. wounds that were ongoing but stable. Biologic functions of these highly expressed proteins were determined by the Gene Ontology Database. In brief, wounds that healed are enriched for proteins involved in skin barrier development, wound healing, blood clot formation, responses to bacteria and external stress. Wounds that deteriorated have higher expression of proteins involved in chronic inflammatory responses, the compliment cascade and a pseudomonas histone kinase. Supplemental figure 3 displays the volcano plot for differential protein expression in younger vs. older wounds.

### Wound Slough is Polymicrobial and Associated with Wound Etiology and Body Site

To assess the microbial bioburden of slough samples, swabs were collected from the wound surface, prior to washing and removal of slough via debridement. Bacterial bioburden was assessed by both quantitative bacterial culture and quantitative-PCR of the bacterial 16S ribosomal RNA gene (Table S5). Slough bioburden was generally high across all samples ranging from 1.0×10^2^ to 8.0×10^7^ colony forming units (CFU) and 4.2×10^3^ to 4.6×10^8^ bacteria per inch^2^ by qPCR. The quantity of bacteria determined by qPCR and the quantity of bacteria detected through quantitative bacterial culture are highly concordant (Fig. 3A, spearman r = 0.84, p-value < 0.01).

**Figure 3:**
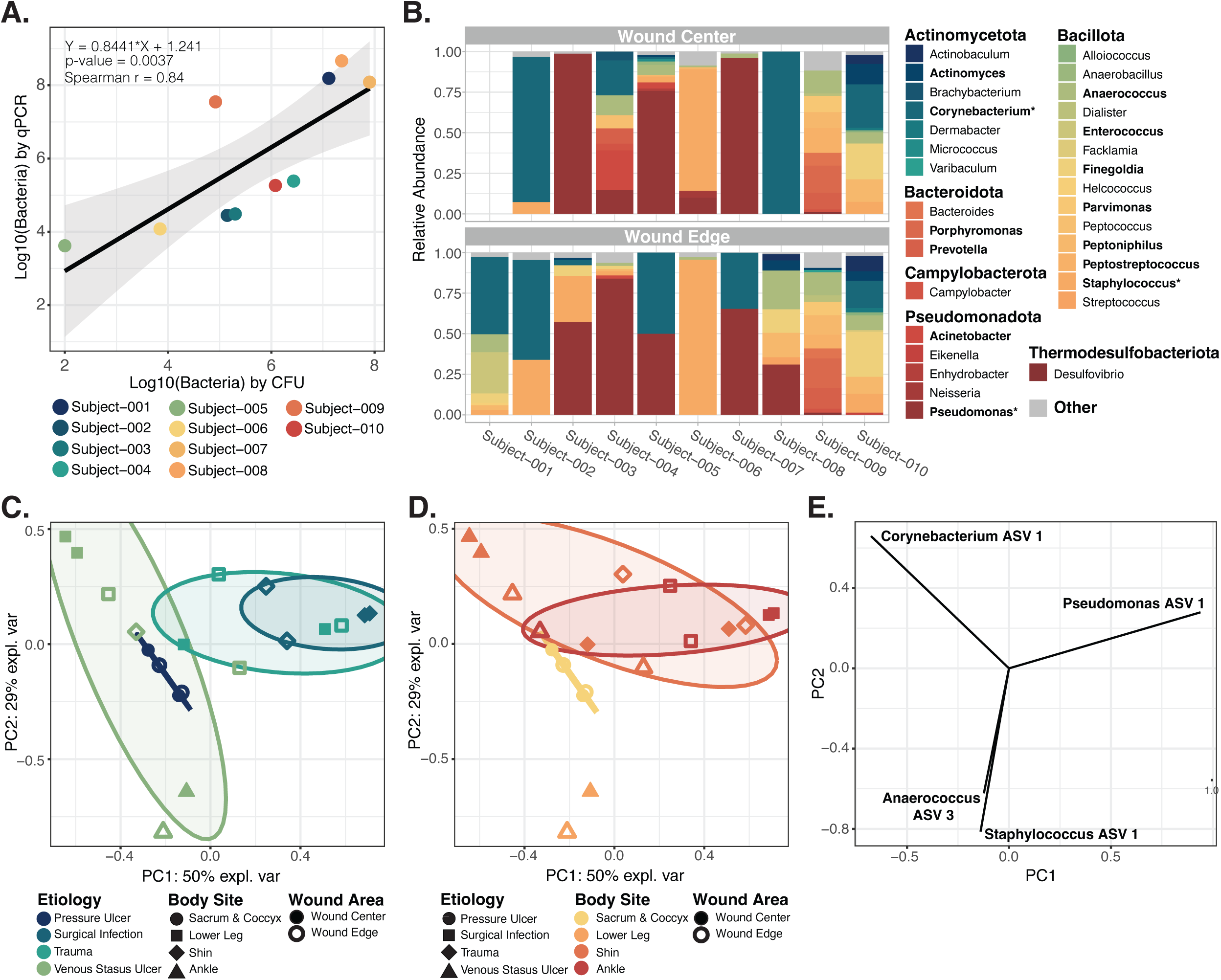
Microbial communities at a wound surface are largely dictated by the body site where the wound is located and the wound’s etiology. Swabs of the surface microbiome were collected from the wound edge and wound center. Subject-001 did not have a sample collected from the wound center due to pain. A) The number of bacteria per inch^2^ determined by quantitative-PCR (qPCR) strongly corelates with the number of bacteria detected through quantitative bacterial culture (measured in bacterial colony forming units [CFU]). Points are colored by subject. B) Relative abundance of bacterial genera on the surface of each subject’s wound center (top) and wound edge(bottom) based on high-throughput sequencing of the bacterial 16S ribosomal gene. For each sample, bacterial taxa that were < 1% abundant were grouped into the “Other” category along with any un-classified bacterial sequences. Genera are grouped by the phyla in which they belong; Actinomycetota (blues), Bacillota (greens to orange-yellows), Bacteroidota (oranges), Campylobacter (deep orange-red), Pseudomonadota (reds), and Thermodesulfobacteriota (deep red). Relative abundance of genera within bolded indicates that the genera comprises > 10% of at least one sample. An * indicates that the genera comprises > 30% of at least one sample. C-D) Principal component analysis indicated that wound surface microbial communities cluster by both the wound’s etiology (C) and the wounds location on the body (D). Plots C and D are the same but colored differently to highlight the sample groupings by etiology and body site respectively. Microbiome samples did not cluster by whether the swab was taken from the wound edge or center, the wounds age, or the wounds outcome 3 months after the sample collection (e.g whether the wound went on to heal or did not). E) A vector plot indicating the primary bacterial ASVs that dictated a points position in the PCA plot C-D. These ASV’s belong to Corynebacterium, Pseudomonas, Staphylococcus, and Anaerococcus species.

To determine the composition and spatial variation of bacterial communities within slough, samples collected from slough at the edge and center of the wound were assessed through high-throughput sequencing of the bacterial 16S ribosomal RNA marker gene (Fig. 3B, Fig. S4). Due to pain, subject-001 did not have a sample collected from the wound center for this analysis. The major bacterial genera detected were consistent with previous wound microbiome studies. Collectively, the most abundant taxa from wound samples include *Corynebacterium* spp., *Pseudomonas* spp., and *Staphylococcus* spp.(Fig. 3B, Fig. S4). Overall microbiome community structure was generally consistent between the wound edge and wound center. However, in some cases microbiome composition drastically differed, such as in subject-008, where a single species appears to dominate the wound center while the wound edge harbors a much more diverse microbiome. Microbial communities dominated by few taxa within the center of the wound more often occurred in the subjects with large (surface area > 25 cm^2^), deep (> 10 cm^3^) wounds.

Principle component analysis was conducted to reduce the dimensionality of the microbiome data set and explore the variability of samples (Fig. 3C-E). For this analysis any bacterial amplicon sequence variants (ASVs) present in only one sample or averaged less than 1% of across all samples were removed. Factors significantly associated with microbial community composition included the wound’s etiology and its location on the body (Type II permutation MANOVA r^2^ =0.47 and = 0.46 respectively, both p-values < 0.01; Fig. 3C-D, Table S6). Notably, community composition was not associated with spatial sampling at the wound edge or center, or the outcome of the wound 3 months following sample collection (p-values > 0.05). The primary bacterial taxa that influenced sample position in the PCA plot belong *to Corynebacterium, Pseudomonas, Staphylococcus,* and *Anaerococcus* species (Fig. 3E).

### Detection of microbial aggregates in slough is highly variable

To evaluate potential commonalities in the microscopic structure of slough and associated microbial aggregates, slough samples were visualized using both confocal scanning laser microscopy (CLSM) and scanning electron microscopy (SEM). Overall, both techniques revealed slough to be highly variable in structure and composition. CLSM of slough histological sections revealed heterogenous, auto-fluorescent fibrinous tissue and DNA (Fig 4 and S5). SEM showed complex milli-, micron-, and nano-meter scale features on the slough surface, consistent with fibrin and collagen fibrils, fibers, and bundles (Fig S6). One semifluid sample (Subject-008) contained undefined crystalline structures.

**Figure 4:**
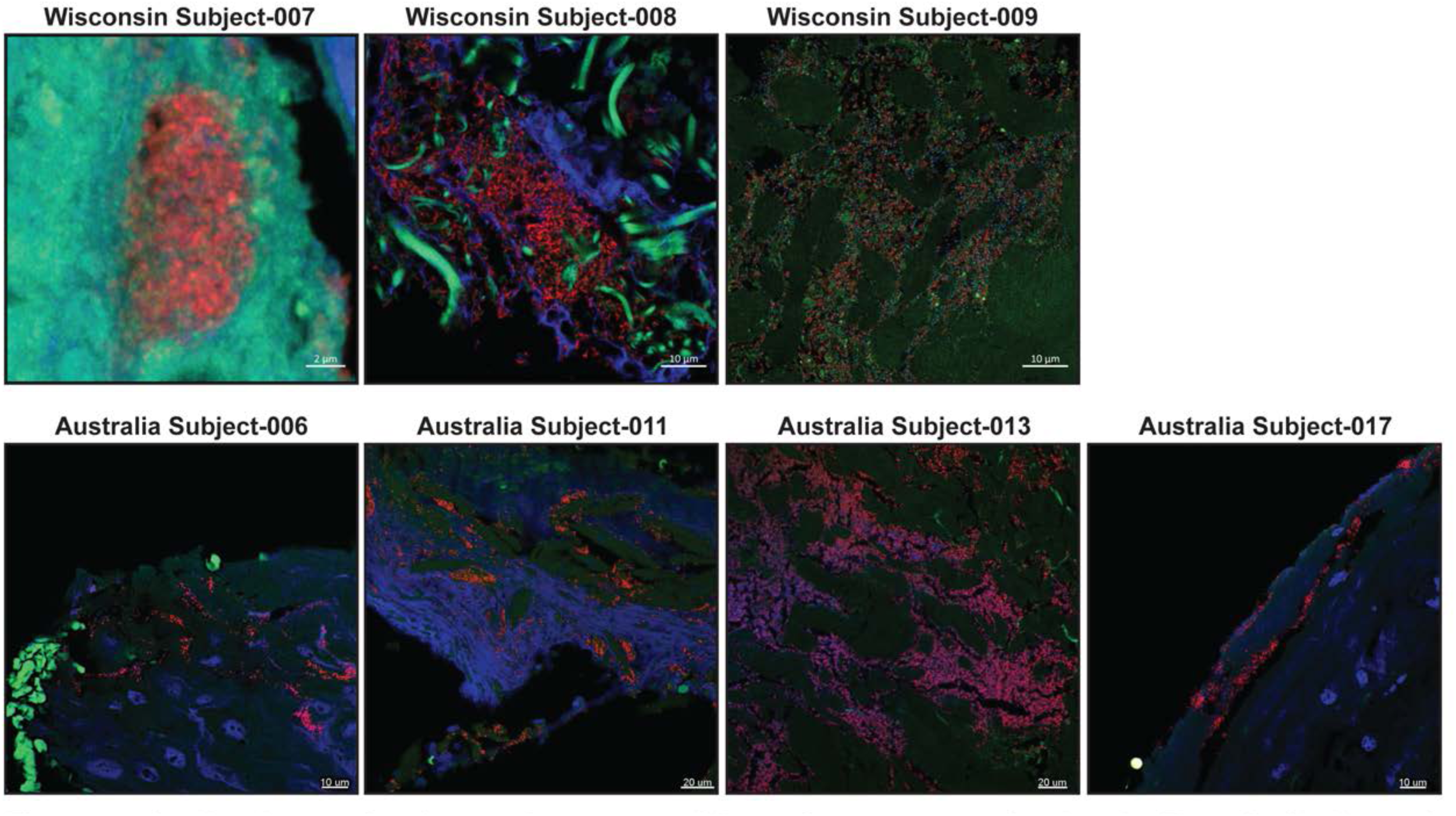
Confocal scanning laser microscopy of bacteria aggregates in slough. Formalin-fixed, paraffin-embedded (FFPE) slough samples were stained with a universal bacterial 16S rRNA probe (red) and for double stranded DNA (DAPI, blue) then visualized with confocal scanning laser microscopy (CSLM). Autofluorescence of the surrounding tissue was visualized in green. Only specimens with detected bacterial aggregates are shown here. Images from specimens with no bacterial aggregates are in supplemental figure 5.

Notably, microbes were only visible in three samples from the Wisconsin cohort, those from patients with the highest slough bioburden, and four of the Australian subjects (Fig 4, S5). This indicates a lower sensitivity of microscopy-based methods. Additionally, all three samples demonstrated different spatial distributions of microbes. Subject-007 had small (∼10μm) aggregates embedded in tissue localized to a DNA-rich, layered region of solid slough (CLSM; Fig 4, S5). Subject-008 had large (>50μm) bacterial aggregates surrounded by extracellular DNA and putative collage fibers within the core of the semifluid slough, suggesting a biofilm community structure (CLSM; Fig 4). Subject-009 had putative collagen bundles colonized with individual rods, cocci, and lancet-shaped bacteria (SEM; Fig S6). CLSM cross-sections showed sparse bacteria in between tissue bundles (Fig 4).

### Integrated analysis reveals key features of non-healing wounds

To predict the variables associated with wound healing outcome, an integrative analysis was pursued encompassing protein clusters, microbial taxa relative abundance, and the numerical Bates-Jensen Wound Assessment score. Datasets were integrated into a supervised Partial Least Squares – Discriminant Analysis (PLS-DA).^34^ To reduce the complexity of the proteomics dataset, K-means clustering was first performed. Further, the top 15 microbial ASVs with greater than 1% relative abundance in at least two subject samples were included (Table S7). PLS-DA revealed that the proteomic and microbial composition of slough and Bates-Jensen scores can distinguish chronic wounds that go on to heal versus those that deteriorate (Fig. 5A,B). Figure 4C illustrates the key variables that help distinguish each outcome group along variate 1 of the PLS-DA plots. Wounds that deteriorated were associated with a higher total Bates-Jensen Assessment score and sub-scores (e.g. higher granulation tissue score, indicating smaller area of the wound bed covered by granulation tissue and poor vascular supply; higher wound edge score, indicating more well-defined to thickened wound edge; as well as greater wound depth); increased abundance of anaerobic taxa (e.g. *Finegoldia* ASV1, *Peptoniphilus* ASV 2), and higher expression of protein clusters 6, 21, and 11. GO Enrichment Analysis revealed that these clusters were enriched for proteins involved in immune responses, particularly immune activation and responses to stimuli, cell motility, and intracellular processes (Fig. 6A). Conversely, wounds that went on to heal were associated with higher abundance of *Acinetobacter* ASV 1 and protein clusters 22, 19, and 5 (Fig. 5C). These clusters were enriched for proteins involved in metabolic and biosynthetic processing, gene expression, and regulation (including negative regulation) of wound healing and responses to stress (Fig. 6C, Fig. S7, Table S8). Overall, the findings of this integrative analysis highlight potentially fundamental differences in the microbial and proteomic composition of slough from wounds that go on to heal compared to those at higher risk for progression.

**Figure 5:**
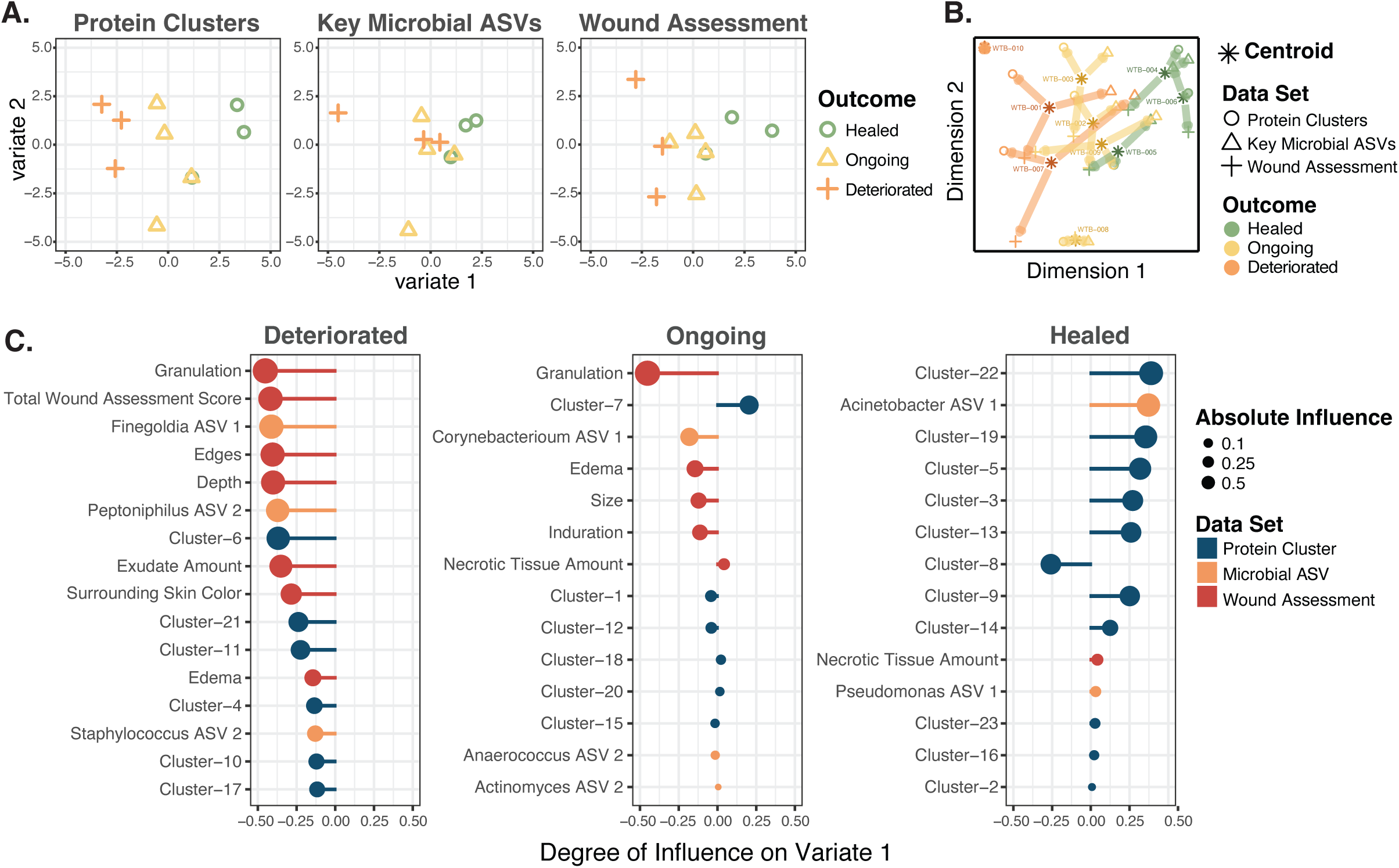
Chronic wounds that go on to heal can be distinguished from those that deteriorate via the proteomic, microbial, and clinical features of slough. To predict the variables associated with wound healing, the protein cluster, microbial, and the Bates – Jensen Wound Assessment datasets were integrated into a supervised Partial Least Squares – Discriminant Analysis (PLS-DA) via the MixOmics package. To simplify the proteomics dataset, proteins were grouped into 23 k-means clusters via the gap-stat method (Table S2). Since there was no significant difference in the microbial community composition at the wound edge or center, samples were combined to create a summative wound slough microbiome for each subject. The “key” 14 microbial ASVs with greater than 1% relative abundance in at least two subjects’ slough samples were included in this integrative analysis (Table S7). A) PLS-DA plots for the protein cluster, microbial ASV, and wound assessment data set respectively. Each dataset contains variables that can distinguish chronic wounds that go on to heal from those that deteriorate. Outcome groups most clearly separate along variate 1 for each of the datasets. B) PLS-DA plot for all the data sets combined. The asterisk indicates the centroid position where the subject’s slough sample falls considering variables from all three datasets. Arrows from the centroid indicate the direction that variables from each individual dataset pull the subject’s datapoint. C)Variable plots of the protein clusters (blue), microbial taxa (orange) and wound assessment criteria (red) that distinguish each outcome group along variate 1 of the PLS-DA plots. A longer vector to the right indicates a variable with greater influence pulling samples to the right along the variate 1 axis. The enriched GO biologic processes for representative slough protein clusters that distinguish slough from wounds with each outcome are in figure 5.

**Figure 6:**
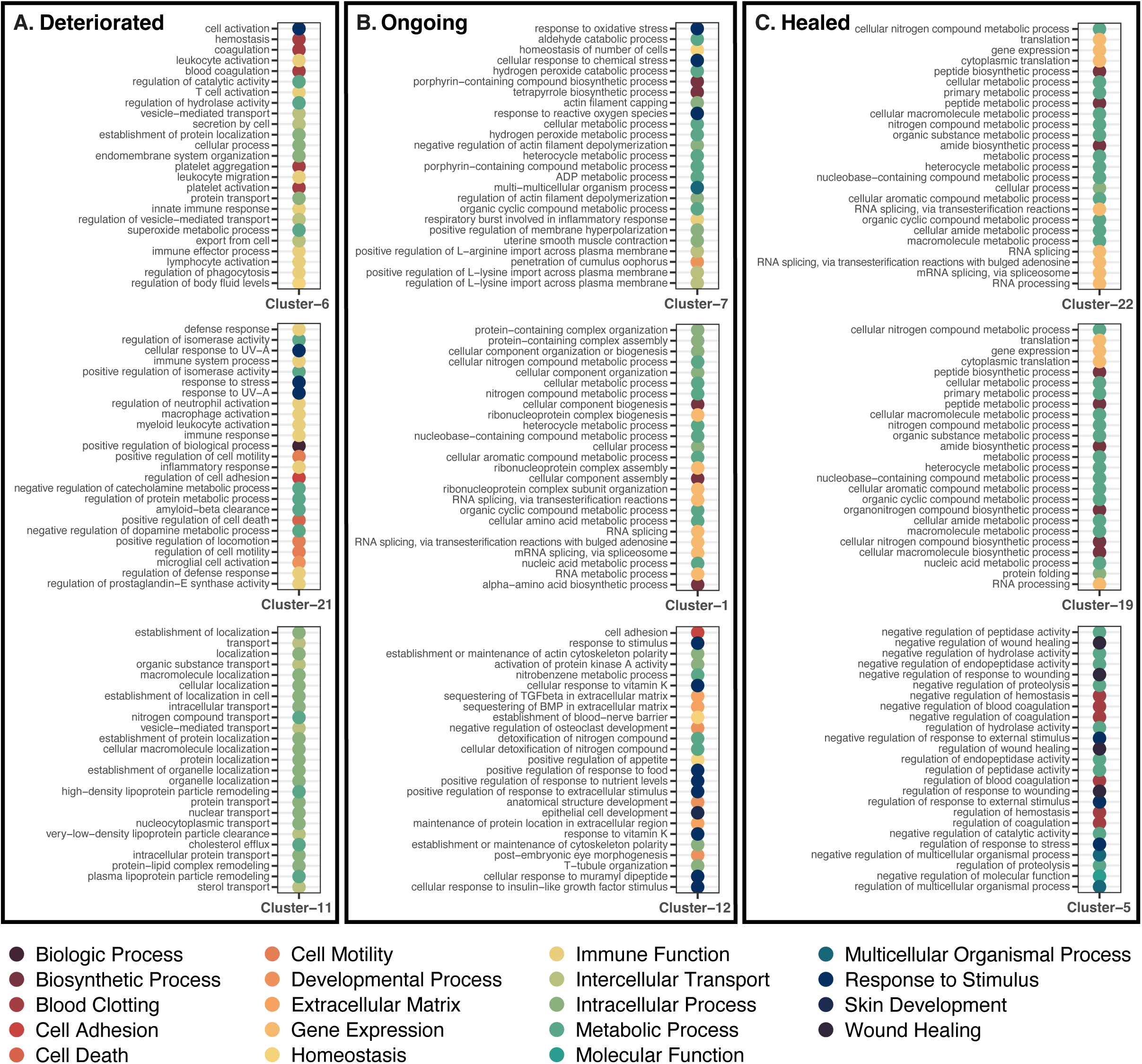
Slough from wounds that go on to deteriorate are enriched for immune activation and inflammatory immune responses. To determine the key biologic processes associated with each protein k-means cluster, proteins within each cluster were submitted as unranked lists to the GO Enrichment analysis tool for evaluation with the PANTHER Overrepresentation test. Details for this analysis are included in supplemental table 8. This figure displays the top 25 most enriched GO biologic process for three representative protein clusters that distinguish slough from wounds that deteriorated three months following sample collection from those that (Fig. 4). For each cluster the biologic processes are ordered from most significantly enriched at the top to least enriched at the bottom. Color of the point indicates the broader biologic classification. A) Wounds that deteriorated are enriched for immune cell activation and inflammatory immune responses, responses to stimuli and stress, cell motility, intracellular transport and intracellular processes. B) Wounds that were stable but ongoing were enriched for responses to stress, metabolic processes, and gene expression. C) Wounds that went on to heal were enriched for metabolic and biosynthetic processing, gene expression, and regulation (particularly negative regulation) of wound healing and responses to stress. Supplemental figure 5 displays the significantly enriched GO biologic processes for all 23 k-means clusters.

## Discussion

Slough is a highly common and burdensome feature of wounds. However, its definition and composition remain poorly characterized. This pilot study aimed to characterize the host and microbial elements of slough across a variety of wound etiologies. We also sought to identify key factors within slough associated with wound healing trajectories. Our findings demonstrate that, i) the microscopic structure of slough is heterogenous and unique to each wound; ii) across subjects wound slough is composed of proteins involved in the structure and formation of the skin, blood clot formation, and various immune responses; iii) the microbial community composition is diverse and corresponds to the wound’s etiology and location on the body; and iv) the composition of slough is associated with wound healing outcomes. Collectively, these findings underscore how the composition of slough itself may be useful for developing microbial and proteomic biomarkers prognostic of wound healing trajectories.

The clinical presentation of slough is highly variable. Slough can range in color from pale yellow to yellow-green, tan, brown, or black to resemble eschar. It can also range in texture from mucous-wet to thick and fibrinous, and range from loosely to firmly attached^6,8^ As expected with this variable clinical presentation, confocal and scanning electron microscopic imaging reveals slough to be microscopically heterogenous and different across wounds (Fig. S5-6). However, as noted by others, slough’s intrinsic gelatinousness consistency makes it easy to perturb and difficult to fix for microscopic assessment.^36^ This likely limits our ability to ascertain additional three-dimensional features within slough that may be pertinent to the wound surface environment.

Microbial biofilm is thought to be highly integrated within wound slough.^21^ In the clinical setting wound slough is often mistaken for microbial biofilm.^15,21^ To address this, several studies have proposed wider adoption of culture based, molecular (i.e. quantitative-PCR), and microscopic techniques into diagnostic practice.^37–40^ At the time of sample collection, only one of the ten subjects was diagnosed with a current wound infection and five had a history of infection in the sampled wound (Table 1). However, SEM and CLSM imaging detected microbes in only three of nine samples tested (Fig. S5-6). Interestingly, subject-009, who had no record of current or previous wound infection, was the only subject to have microbes visualized via both SEM and CLSM. This speaks to the difficulty in identifying biofilm or even the presence of single cells of bacteria using microscopy techniques as more sensitive molecular methods indicated every sample contained a considerable bioburden of bacteria. Further, the detection rates for microbial biofilm in this study are notably lower than previously reported for chronic wound samples.^39–41^ This could be due to a number of factors, including spatial heterogeneity of bacterial aggregates across the wound surface. Indeed, to saturate sampling efforts hundreds of slides and images would need to be obtained. To improve detection rates the incorporation of methods that increase specificity of bacterial detection such as immunogold labeling or gold in situ hybridization could be applied, but remain impractical for routine clinical evaluation.^42,43^

The quantity of bacteria determined via qPCR correlates with the bacterial burden as determined by quantitative culture (Fig. 3A).The reference standard for clinical definition of a wound infection is 10^5^ or more cultured colony forming units (CFU)/ml.^37^ By that metric, eight of the ten subjects meet definition for clinical infection, despite an absence of clinical sign of infection (Fig. 3A). Indeed, only one subject had a diagnosed infection. While the use of qPCR for detecting bacterial bioburden is more sensitive, particularly for patients like subject-009 whose wounds may contain more anaerobic or difficult to culture bacteria (Fig. 3A,B), this data suggests the use of such cutoffs are complicated and should be used with caution. Indeed, wounds with high bacterial bioburden can go on to heal without intervention with antibiotics.

Isolation and identification of bacteria from all subjects through both microbial culture and 16S sequencing underscores that even in the absence of a clinical wound infection, slough contains complex microbial communities (Fig. 3A-B, Table S5). Previous work evaluating the influence of sharp debridement on the wound microbiome further has shown that these microbes are likely highly integrated within and throughout wound slough.^16^ Here, the most frequently isolated via microbial culture were *Corynebacterium, Staphylococcus,* and *Pseudomonas* species (Table S6). *Corynebacterium, Staphylococcus,* and *Pseudomonas* were also the most abundant taxa identified via 16S profiling, comprising at least 30% of the microbial community in slough three of the ten subjects respectively (Fig. 3B). Across wound etiologies *Corynebacterium, Staphylococcus,* and *Pseudomonas* species appear to be the most abundant taxa within the chronic wound microbiome (Fig. 3B).^16–18,44^ Contradicting some previous reports,^16,18,44^ we find slough microbial community composition to be associated wound etiology and location on the body (Fig. 3C-E, Table S6). The microbiome of healthy intact skin naturally varies across body sites due to differences in the physiologic characteristics of the local skin environment.^45–49^ It is plausible that these variations in the microbiome of the surrounding skin influence the community structures within wound slough.

In terms of the human components of the wound, there are only a handful of reports on the proteomic composition of tissue biopsies, granulation tissue, and exudative fluid from chronic pressure ulcers and diabetic wounds.^9–14^ Broadly speaking, these studies find fluid and tissue from these wounds to contain various types of collagen, extracellular matrix proteins, matrix metalloproteases, clotting factors and proteins generally related to innate and acute immune responses.^9–14^ To our knowledge, this is the first study to specifically evaluate the proteomic composition of wound slough. In line with these previous studies, slough from chronic wounds is primarily composed of keratin and various types of collagen, extracellular matrix proteins, matrix metalloproteases, clotting factors, and immune response proteins (Table S2, Fig. 1). More specifically, slough is enriched for proteins involved in skin barrier integrity and formation, wound healing, and immune functions ranging from innate compliment activation to acute responses to stimuli (e.g. to bacteria) and humoral immune responses (Fig. 1, Fig. S2). The high prevalence of intracellular and skin associated proteins combined with the relative absence of enrichment for vascular and angiogenic pathways supports that hypothesis that slough is largely devitalized tissue. However, many of these proteins may be functional in this environment. Further, the collective abundance of proteins associated with inflammatory cells such as neutrophils, underscores the leading theory that slough is a biproduct of prolonged inflammatory process.^6,7^

Of the three main data sets assessed (proteomics, microbiome, and the Bates Jensen Wound Assessment), the proteomics dataset had the strongest associations with wound healing outcome. When assessed independently, Bates-Jensen Wound Assessment Tool (BWAT) scores were not significant for wounds that deteriorated, nor were there associations between microbiome composition and wound outcome (Table 2, Fig. 3). Although these analyses were likely limited due to low subject numbers, differential Protein Expression Analysis (DeqMS)^30^ found wounds that went on to heal were enriched for proteins involved in skin barrier development, wound healing, blood clot formation, and responses to bacteria and external stress (Fig. 2B-D, Table S4). Conversely, deteriorating wounds were enriched for proteins involved in immune responses categorized as chronic inflammatory responses and the compliment cascade. Of the proteins enriched in wounds that deteriorated; AP-1 is a notable biphasic regulator of wound healing^50^; NLR family proteins and Caveolase-associated protein 1 have been associated with impaired wound healing in murine models^51–53^; and CD177, compliment factor H, and vasodilator-stimulated phosphoprotein have also been noted to be elevated in chronic wound fluid and/or tissue.^11,12^ To identify variables associated with wound healing we incorporated proteomics, microbiome, and the BWAT score datasets into a supervised Partial Least Squares – Discriminant Analysis (PLS-DA).^34^ In this model, slough from wounds that healed were enriched for proteins involved in regulation, particularly negative regulation, of immune responses and wound healing as well as the aerobic microbial taxa *Acinetobacter* (Fig. 5-6). Conversely, wounds that deteriorated contained slough enriched with inflammatory proteins, particularly those involved in immune activation, responding to stimuli and chronic inflammation. Wounds that deteriorated were also associated with greater abundance of anaerobic microbial taxa, *Finegoldia* and *Peptoniphilus*, as well as higher Bates-Jensen wound assessment scores, indicating a more severe wound state.

Overall, our model’s findings are consistent with related literature. For instance, they underscore BWAT’s clinical utility in a well-rounded wound evaluation, and suggest that high sub-scores for granulation, wound edge, wound depth, and exudate amount may hold the strongest predictive potential for identifying a wound likely to deteriorate (Fig. 5).^24,35,54^ From a microbiological perspective, high abundance of anaerobic taxa and select *Staphylococci* species are frequently associated with impaired wound healing and poor outcomes.^16,17,55,56^ Similar proteomic investigations with wound fluid and tissue biopsies also find elevated inflammatory proteins and enrichment of proteases and matrix metalloproteinases in wounds that do poorly as well as enrichment of extracellular matrix proteins and keratin in healing wounds.^10,12^ This work demonstrates that slough, which is often regularly debrided as a part of standard care, provides a readily available, underutilized, high protein concentration biomarker reservoir. Most of the proteins identified through independent DeqMS assessment also fall within the protein clusters that distinguish between healed and non-healing wounds in the comprehensive model (Fig. 2 and 5, Tables S2, S4 and S8), suggesting these identified proteins have the greatest potential as biomarker targets to predict wound healing.

The primary limitation of this pilot study is the small number of samples enrolled. Future investigations intend to expand upon these methods, potentially with even more targeted proteomic and microbiologic approaches, to validate the predicted features associated with wound healing outcome in a larger cohort. This study is also limited in the collection of tissue samples from the wound bed itself, after removal of slough. Future studies should consider collecting both slough and wound tissue samples to understand the proportion of slough proteins that overlap with proteins also found in the wound bed. There were also several factors that inhibited the microscopic detection of microorganisms via SEM and CLSM. For instance, initial cleansing of the wound with soap and water prior to debridement per standard of care, may have removed superficial microbial aggerates. With SEM, there are no efficient algorithms for distinguishing individual microbes or microbial biofilm from background collagen fibers and tissue. The heterogeneous and often gelatinous texture of slough along with the ability to only view a histological cross-section, may have also limited microbial detection via CLSM. Our microbial assessment with 16S amplicon sequencing only provides genus level resolution, and not all species within a genus have propensity to cause infection (e.g. *Staphylococcus aureus* vs. *Staph. hominis).* Evaluating chronic wound metagenomes would provide species level resolution and detect the presence of virulence and antibiotic resistance genes. However, to date there are very few investigations into wound microbial metagenomes as this method remains limited due to cost.^17^

In conclusion, slough is an underutilized reservoir for potential microbial and proteomic biomarkers. To our knowledge this is the first study to integrate clinical wound assessment, microbiome, and proteomic data into a single assessment for the prediction of wound healing outcome. Future studies intend to utilize these and similar methods to further explore the biomarkers within slough in a larger cohort with appropriate statistical power. Utilization of a comprehensive patient-centered assessment will lead to more effective identification of high-risk patients wounds for triage into specialty care, ultimately, reducing the healthcare, financial, and personal burden of living with hard to heal wound.

## Acknowledgements

Special thanks to Derek A. Gonzalez and the UW-Heath Department of Surgery Clinical Research Team for assisting with subject recruitment, enrollment, and sample collection.

We also like to thank the UW-Biotechnology Center for microbial sequencing, mass-spectrometry, and initial proteomics analysis.

## Contributions

Conceptualization: L.R.K, G.S, T.S, K.O

Methodology: L.R.K, T.B, M.M, E.C.T, J.Z.C, M.R, B.F

Analysis: L.R.K, E.C.T, A.C, T.B, M.M, M.R, B.F

Investigation: L.R.K, A.G, M.M, T.B, E.C.T, J.Z.C, M.R, B.F

Resources: L.R.K, A.G, M.M, T.B

Writing – Original Draft: E.C.T, J.Z.C, L.R.K, A.G

Writing – Review & Editing: E.C.T, J.Z, G.S, M.M, T.B, M.R, B.F, K.O, T.S, A.G, L.R.K

Visualization – E.C.T, A.C, M.R, B.F Supervision – L.R.K, A.G, T.B, M.M

## Supplemental Figure Legends

**Supplemental Figure 1: Photos of subject wounds before debridement procedure**.

**Supplemental Figure 2: Directed Acyclic Graph (DAG) of the significantly enriched gene ontology (GO) terms grouped by biologic processes (A) molecular functions (B) and cellular components (C) within wound slough.** The most abundant proteins across all slough debridement tissue samples were input as a ranked list to the Gene Ontology enRIchment analysis (GORILA) and visualization tool.^29^ Figure 1 displays the enriched GO terms associated with each DAG. The significantly enriched GO terms for each DAG are displayed. Box colors indicate p-values; white > 10^-3^; yellow 10^-3^ – 10^-5^, yellow-orange 10^-5^ – 10^-7^, orange 10^-7^ – 10^-9^, Red < 10^-9^. More detail, including GO term annotations, descriptions, enrichment, number of proteins (Uniprot Genes) involved from our dataset involved in each GO Term, and FDR-qValues are in Supplemental Table 3.

**Supplemental Figure 3: Chronic wounds present less than 1 year are enriched for proteins involved in epithelial barrier formation, neutrophil degranulation, and response to bacteria.** Conversely, wounds present for more than 1 year are enriched for proteins involved in iron sequestration and tRNA metabolism. Subjects were grouped the age of the wound at the time of sample collection. Wounds present for less than 1 year were considered “young”, and those present for more than 1 year were considered “old.” Groups were assessed for differential protein expression via DEqMS. This volcano plot displays the proteins with significantly greater expression in younger or older wounds.

**Supplemental Figure 4: The Abundance of key bacterial taxa is similar across wound slough from two distinct subject cohorts from Wisconsin and Australia.** Datasets were generated using amplicon sequencing of the V4 (panel A, Wisconsin) or V1V3 (panel B, Australia) regions of the 16S rRNA gene, and were thus analyzed separately. ASVs were summed at the genus level. Note that the taxonomic resolution for classification may differ by amplicon region. Genera are shown if present at above 5% relative abundance in at least one specimen and are ordered by mean relative abundance across all specimens within a dataset. Subjects are ordered by average linkage hierarchical clustering of Bray-Curtis dissimilarities. In the Wisconsin cohort (panel A), subject taxa profiles are averaged from multiple specimens.

**Supplemental Figure 5: Confocal scanning laser microscopy of slough samples without bacterial aggregates.** Formalin-fixed, paraffin-embedded (FFPE) slough samples were stained with a universal bacterial 16S rRNA probe (red) and for double stranded DNA (DAPI, blue) then visualized with confocal scanning laser microscopy (CSLM). Autofluorescence of the surrounding tissue was visualized in green. The specimens with detected bacterial aggregates are shown in figure 4. Here are the remaining specimens from both patient cohorts that did not have identifiable bacterial aggregates.

**Supplemental Figure 6: Scanning electron microscopy finds slough to be variable in structure and unique to the subject.** Debrided slough samples were evaluated via scanning electron microscopy (SEM). Subjects-004, –005, and –006 did not have enough debridement tissue for SEM. One subject, subject-009 had visible microorganisms on SEM. A majority of specimens were fibrous in appearance, while one specimen had crystalline structures.

**Supplemental Figure 7: Enriched GO biologic processes for each of the 23 k-means protein clusters.** To determine the key biologic processes associated with each protein k-means cluster, proteins within each cluster were submitted as unranked lists to the GO Enrichment analysis tool for evaluation with the PANTHER Overrepresentation test. Details for this analysis are included in supplemental table 8. This figure displays the top 25 most enriched GO biologic process each of the 23 k-means protein clusters. For each biologic processes are ordered from most significantly enriched at the top to least enriched at the bottom. Color of the point indicates the broader biologic classification.

## Supplemental Table Legends

**Supplemental table 1: Detailed subject and wound characteristics.** Ten subjects with chronic or slow to heal wounds of various etiologies were enrolled from the UW-Health Wound Care Clinic. Wounds were evaluated with the Bates-Jensen Wound Assessment Tool. Information on the wound and patient comorbidities were extracted from the medical record at the time of sample collection. Information on whether the wound went on to heal, was ongoing yet clinically stable, or deteriorated 3 months following sample collection was also recorded. This table serves as a compliment to table 1 and 2, providing subject level detail on the wound and patient comorbidities.

**Supplemental Table 2: Proteomic composition of wound slough.** Normalized and means centered protein peptide abundance within each subject’s wound slough. Description, species of origin (eg. Homosapiens or bacteria), broad GO biological process, GO cellular component, GO molecular function, WikiPathways, Reactome Pathways, and KEGG pathways for each protein peptide accession are also included. To simplify the proteomics dataset for integrative PLS-DA analysis proteins were grouped into 23 clusters via the gap-stat method. The Kmeans cluster in which the protein falls is also indicated.

**Supplemental Table 3: Data-frame of the GO terms that are significantly enriched in chronic wound slough.** Debrided slough tissue was sent for proteomic characterization via mass spectrometry. A) The most abundant proteins across all samples were input as a ranked list to the Gene Ontology enRIchment analysis (GORILA) and visualization tool.^29^ Significantly enriched GO terms are listed with their GO term annotations, descriptions, enrichment, number of proteins (Uniprot Genes) involved from our dataset involved in each GO Term, and FDR-qValues. Visual representations of this data can be found in figure 1 and supplemental figure 2.

**Supplemental Table 4: Proteins with significantly greater expression subjects grouped by outcome or wound age.** Subjects were grouped by the wound’s outcome (healed, ongoing, deteriorated), or wound age (young [wounds present < 1 year], or old [wounds present > 1 year]) and groups were assessed for differential protein expression via DEqMS. Only proteins with significantly greater expression (log_2_ Fold change > 1 and log_10_ P-value < 10^-2^) are displayed. Details on the protein description and associated GO terms, WikiPathways, Reactome Pathways, and KEGG pathways are included. Volcano plots indicating the proteins with significantly greater expression in each of the associated group comparisons are in figure 2 B-D and supplemental figure 5.

**Supplemental Table 5: Bacterial bioburden and Identification of Cultured bacteria from the wound surface.** Swabs of the wound slough microbiome were collected into either DNA/RNA Shield or liquid ames broth. Bacterial DNA was extracted from the samples collected into DNA/RNA Shield, and bacterial bioburden was assessed through quantitative PCR of the 16S ribosomal gene. Samples collected into liquid ames broth were plated on to blood agar and grown overnight at 37C for quantitative bacterial culture. Individual bacterial colonies with distinct morphologies were isolated and grown overnight. To identify the genus of these isolates, bacterial DNA was extracted and sent for sanger sequencing of the 16S bacterial ribosomal gene. Genera of successfully identified isolates are listed.

**Supplemental Table 6: Univariate type 2 permutation MANOVA results indicate that microbial community composition were the wound’s etiology and its location on the body.** Table of univariate type 2 permutation MANOVA results. Each permutation MANOVA was run via the Euclidian method with 9999 permutations using the Adonis 2 r package. * Indicates p-value less than 0.05. ** Indicates p-value less than 0.01.

**Supplemental Table 7: relative abundance for key microbial ASV’s in wound slough.** Since there was no significant difference in the microbial community composition at the wound edge or center (Type II permutation MANOVA p-value > 0.5, Table S6), samples were combined to create a summative wound slough microbiome for each subject. Relative abundance of each ASV in the wound center and wound edge were averaged. This table displays the 14 microbial ASVs with greater than 1% relative abundance in at least two summative subject slough microbiome samples, which were included in the integrative PLS-SA analysis. Anaerococcus ASV 3 (6a418787996565e7641dbbf39b7d3e18) and Staphylococcus ASV 1 (18af7b7f2b61429936fcd63a453fcefd) from figure 3 are not included here since they were only present in samples from one subject (subject-006) and subsequently not included in the PLS-DA. Although not included in the PLS-DA analysis, the relative abundance of all other taxa, which were either only present in samples from that subject, or present at < 1% is also indicated to highlight the proportion of taxa within a subject that were of low abundance or unique to that subject.

**Supplemental Table 8: Most enriched GO biologic processes for each of the 23 k-means protein clusters.** To determine the key biologic processes associated with each protein k-means cluster, proteins within each cluster were submitted as unranked lists to the GO Enrichment analysis tool for evaluation with the PANTHER Overrepresentation test. This table depicts the 25 most significantly enriched GO biologic processes for each protein cluster and includes the associated go terms, the broader classification, number of protein IDs in *Homo sapiens* reference database, number of IDs in uploaded K-means cluster, the expected number of IDs, fold enrichment, p-value, and false discovery rate (FDR) if it was able to be calculated. To determine the most enriched biologic processes, terms by the smallest to largest FDR, followed by smallest to largest p-value if FDR was unable to be calculated. A rank of 1 indicates that it was the most enriched biologic process in the protein cluster. Supplemental table 2 includes details on the proteins within each cluster.

## Works Cited

1. Nussbaum, S. R. et al. An Economic Evaluation of the Impact, Cost, and Medicare Policy Implications of Chronic Nonhealing Wounds. Value in Health 21, 27–32 (2018).

2. Sen, C. K. Human Wound and Its Burden: Updated 2020 Compendium of Estimates. Advances in Wound Care 10, 281–292 (2021).

3. Guest, J. F., Fuller, G. W. & Vowden, P. Cohort study evaluating the burden of wounds to the UK’s National Health Service in 2017/2018: update from 2012/2013. BMJ Open 10, e045253 (2020).

4. McCosker, L. et al. Chronic wounds in Australia: A systematic review of key epidemiological and clinical parameters. International Wound Journal 16, 84–95 (2019).

5. Evans, K. & Kim, P. J. Overview of treatment of chronic wounds. UpToDate https://www.uptodate.com/contents/overview-of-treatment-of-chronic-wounds#H45052022 (2022).

6. Angel, D. Slough: what does it mean and how can it be managed. WPR 27, (2019).

7. Grey, J. E., Enoch, S. & Harding, K. G. Wound assessment. BMJ 332, 285–288 (2006).

8. McGuire, J. & Nasser, J. J. Redefining Slough: A New Classification System to Improve Wound Bed Assessment and Management. Wounds 3, 61–66 (2021).

9. Jia, Z. et al. Proteomics changes after negative pressure wound therapy in diabetic foot ulcers. Mol Med Rep 24, 834 (2021).

10. Eming, S. A. et al. Differential Proteomic Analysis Distinguishes Tissue Repair Biomarker Signatures in Wound Exudates Obtained from Normal Healing and Chronic Wounds. J. Proteome Res. 9, 4758–4766 (2010).

11. Edsberg, L. E., Wyffels, J. T., Brogan, M. S. & Fries, K. M. Analysis of the proteomic profile of chronic pressure ulcers: Proteomics of chronic pressure ulcers. Wound Repair Regen 20, 378–401 (2012).

12. Baldan-Martin, M. et al. Comprehensive Proteomic Profiling of Pressure Ulcers in Patients with Spinal Cord Injury Identifies a Specific Protein Pattern of Pathology. Advances in Wound Care 9, 277–294 (2020).

13. Krisp, C. et al. Proteome analysis reveals antiangiogenic environments in chronic wounds of diabetes mellitus type 2 patients. Proteomics 13, 2670–2681 (2013).

14. Fadini, G. P. et al. The molecular signature of impaired diabetic wound healing identifies serpinB3 as a healing biomarker. Diabetologia 57, 1947–1956 (2014).

15. Schultz, G. et al. Consensus guidelines for the identification and treatment of biofilms in chronic nonhealing wounds: Guidelines for chronic wound biofilms. Wound Rep and Reg 25, 744–757 (2017).

16. Verbanic, S., Shen, Y., Lee, J., Deacon, J. M. & Chen, I. A. Microbial predictors of healing and short-term effect of debridement on the microbiome of chronic wounds. npj Biofilms Microbiomes 6, 21 (2020).

17. Kalan, L. R. et al. Strain– and Species-Level Variation in the Microbiome of Diabetic Wounds Is Associated with Clinical Outcomes and Therapeutic Efficacy. Cell Host & Microbe 25, 641-655.e5 (2019).

18. Wolcott, R. D. et al. Analysis of the chronic wound microbiota of 2,963 patients by 16S rDNA pyrosequencing. Wound Rep and Reg 24, 163–174 (2016).

19. Kvich, L., Burmølle, M., Bjarnsholt, T. & Lichtenberg, M. Do Mixed-Species Biofilms Dominate in Chronic Infections?–Need for in situ Visualization of Bacterial Organization. Front. Cell. Infect. Microbiol. 10, 396 (2020).

20. Lichtenberg, M. et al. Single cells and bacterial biofilm populations in chronic wound infections. APMIS apm.13344 (2023) doi:10.1111/apm.13344.

21. Percival, S. L. & Suleman, L. Slough and biofilm: removal of barriers to wound healing by desloughing. J Wound Care 24, 498–510 (2015).

22. Powers, J. G., Higham, C., Broussard, K. & Phillips, T. J. Wound healing and treating wounds. Journal of the American Academy of Dermatology 74, 607–625 (2016).

23. Frykberg, R. G. & Banks, J. Challenges in the Treatment of Chronic Wounds. Advances in Wound Care 4, 560–582 (2015).

24. Bates-Jensen, B. Bates-Jensen Wound Assessment Tool. (2001).

25. Malone, M., Radzieta, M., Schwarzer, S., Jensen, S. O. & Lavery, L. A. Efficacy of a topical concentrated surfactant gel on microbial communities in non-healing diabetic foot ulcers with chronic biofilm infections: A proof-of-concept study. International Wound Journal 18, 457–466 (2021).

26. Loesche, M. et al. Temporal Stability in Chronic Wound Microbiota Is Associated With Poor Healing. J. Invest. Dermatol. 137, 237–244 (2017).

27. Bolyen, E. et al. Reproducible, interactive, scalable and extensible microbiome data science using QIIME 2. Nat Biotechnol 37, 852–857 (2019).

28. McMurdie, P. J. & Holmes, S. phyloseq: An R Package for Reproducible Interactive Analysis and Graphics of Microbiome Census Data. PLoS ONE 8, e61217 (2013).

29. Eden, E., Navon, R., Steinfeld, I., Lipson, D. & Yakhini, Z. GOrilla: a tool for discovery and visualization of enriched GO terms in ranked gene lists. BMC Bioinformatics 10, 48 (2009).

30. Zhu, Y. et al. DEqMS: A Method for Accurate Variance Estimation in Differential Protein Expression Analysis. Molecular & Cellular Proteomics 19, 1047–1057 (2020).

31. Mi, H., Muruganujan, A., Casagrande, J. T. & Thomas, P. D. Large-scale gene function analysis with the PANTHER classification system. Nat Protoc 8, 1551–1566 (2013).

32. Mi, H., Muruganujan, A., Ebert, D., Huang, X. & Thomas, P. D. PANTHER version 14: more genomes, a new PANTHER GO-slim and improvements in enrichment analysis tools. Nucleic Acids Research 47, D419–D426 (2019).

33. Nadler, N. et al. The discovery of bacterial biofilm in patients with muscle invasive bladder cancer. APMIS 129, 265–270 (2021).

34. Rohart, F., Gautier, B., Singh, A. & Lê Cao, K.-A. mixOmics: An R package for ‘omics feature selection and multiple data integration. PLoS Comput Biol 13, e1005752 (2017).

35. Harris, C. et al. Bates-Jensen Wound Assessment Tool: Pictorial Guide Validation Project. *Journal of Wound*, Ostomy and Continence Nursing 37, 253–259 (2010).

36. Lange-Asschenfeldt, S. et al. Applicability of confocal laser scanning microscopy for evaluation and monitoring of cutaneous wound healing. J. Biomed. Opt. 17, 1 (2012).

37. Tuttle, M. S. Association Between Microbial Bioburden and Healing Outcomes in Venous Leg Ulcers: A Review of the Evidence. Advances in Wound Care 4, 1–11 (2015).

38. Li, S., Renick, P., Senkowsky, J., Nair, A. & Tang, L. Diagnostics for Wound Infections. Advances in Wound Care 10, 317–327 (2021).

39. Oates, A. et al. The Visualization of Biofilms in Chronic Diabetic Foot Wounds Using Routine Diagnostic Microscopy Methods. Journal of Diabetes Research 2014, 1–8 (2014).

40. Hurlow, J., Blanz, E. & Gaddy, J. A. Clinical investigation of biofilm in non-healing wounds by high resolution microscopy techniques. J Wound Care 25, S11–S22 (2016).

41. Johani, K. et al. Microscopy visualisation confirms multi-species biofilms are ubiquitous in diabetic foot ulcers: Biofilms in DFUs. Int Wound J 14, 1160–1169 (2017).

42. Davis, C. L. & Brlansky, R. H. Use of Immunogold Labelling with Scanning Electron Microscopy To Identify Phytopathogenic Bacteria on Leaf Surfaces. Appl Environ Microbiol 57, 3052–3055 (1991).

43. Ye, J., Nielsen, S., Joseph, S. & Thomas, T. High-Resolution and Specific Detection of Bacteria on Complex Surfaces Using Nanoparticle Probes and Electron Microscopy. PLoS ONE 10, e0126404 (2015).

44. Mahnic, A., Breznik, V., Bombek Ihan, M. & Rupnik, M. Comparison Between Cultivation and Sequencing Based Approaches for Microbiota Analysis in Swabs and Biopsies of Chronic Wounds. Front. Med. 8, 607255 (2021).

45. Grice, E. A. & Segre, J. A. The skin microbiome. Nature Reviews Microbiology 9, 244– 253 (2011).

46. Costello, E. K. et al. Bacterial Community Variation in Human Body Habitats Across Space and Time. Science 326, 1694–1697 (2009).

47. Oh, J. et al. Biogeography and individuality shape function in the human skin metagenome. Nature 514, 59–64 (2014).

48. Townsend, E. C. & Kalan, L. R. The dynamic balance of the skin microbiome across the lifespan. Biochemical Society Transactions 51, 71–86 (2023).

49. Swaney, M. H., Nelsen, A., Sandstrom, S. & Kalan, L. R. Sweat and Sebum Preferences of the Human Skin Microbiota. Microbiol Spectr 11, e04180–22 (2023).

50. Neub, A., Houdek, P., Ohnemus, U., Moll, I. & Brandner, J. M. Biphasic Regulation of AP-1 Subunits during Human Epidermal Wound Healing. Journal of Investigative Dermatology 127, 2453–2462 (2007).

51. Qin, Y. et al. NLRC3 deficiency promotes cutaneous wound healing due to the inhibition of p53 signaling. Biochimica et Biophysica Acta (BBA) – Molecular Basis of Disease 1868, 166518 (2022).

52. Jozic, I. et al. Glucocorticoid-mediated induction of caveolin-1 disrupts cytoskeletal organization, inhibits cell migration and re-epithelialization of non-healing wounds. Commun Biol 4, 757 (2021).

53. Boodhoo, K., Vlok, M., Tabb, D. L., Myburgh, K. H. & Van De Vyver, M. Dysregulated healing responses in diabetic wounds occur in the early stages postinjury. Journal of Molecular Endocrinology 66, 141–155 (2021).

54. Smet, S. et al. The measurement properties of assessment tools for chronic wounds: A systematic review. International Journal of Nursing Studies 121, 103998 (2021).

55. Loesche, M. et al. Temporal Stability in Chronic Wound Microbiota Is Associated With Poor Healing. Journal of Investigative Dermatology 137, 237–244 (2017).

56. Choi, Y. et al. Co-occurrence of Anaerobes in Human Chronic Wounds. Microb Ecol 77, 808–820 (2019).

