## Supplementary material for "What is Slough? A pilot study to define the proteomic and microbial composition of wound slough and its implications for wound healing": Sup. Fig. 1

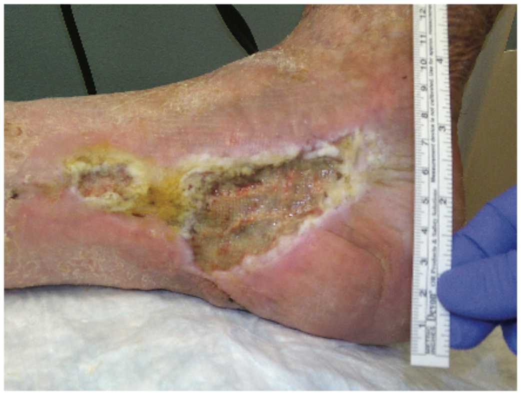

Subject -001

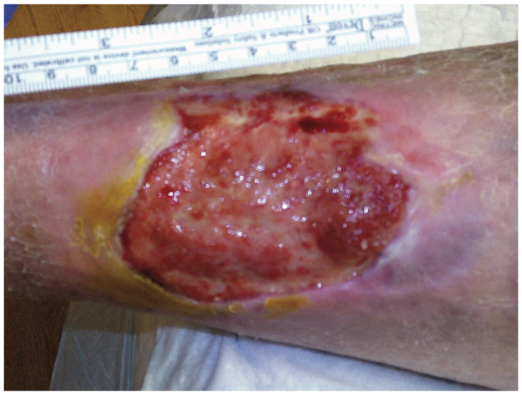

Subject -002

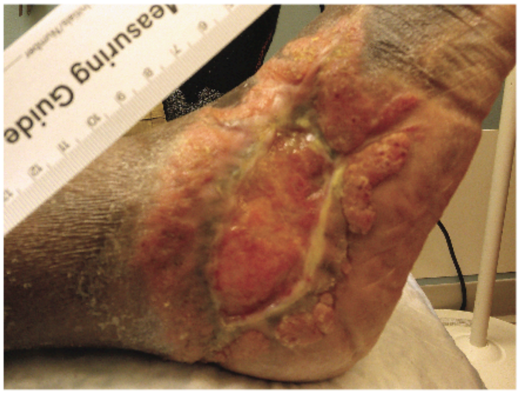


Subject -003

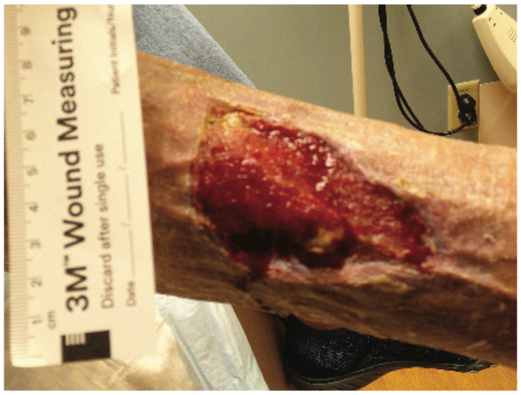


Subject -004

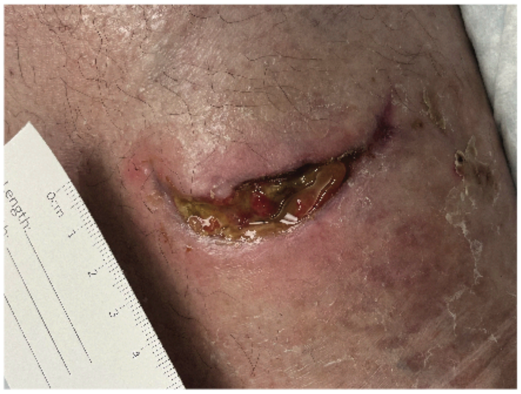

Subject -005

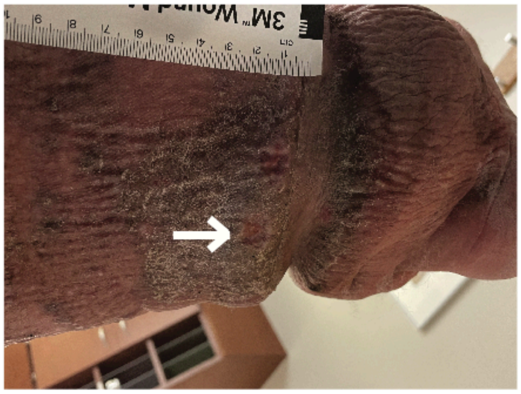

Subject -006

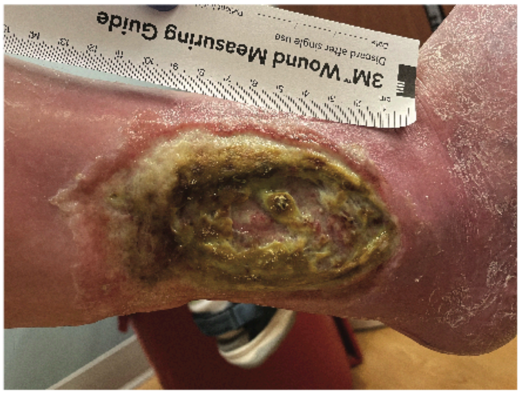


Subject -007

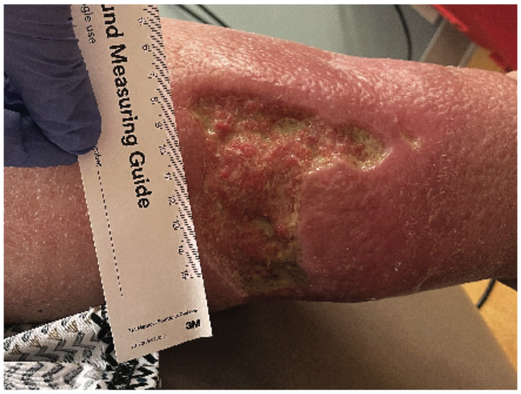


Subject -008

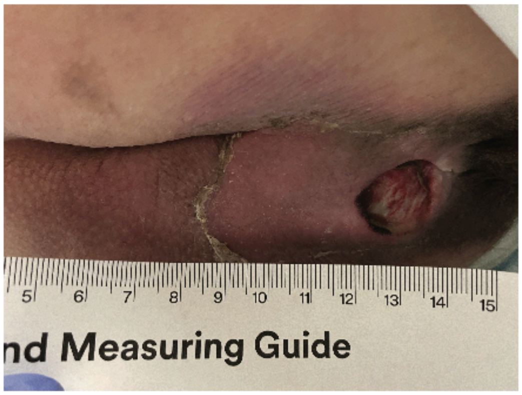


Subject -009

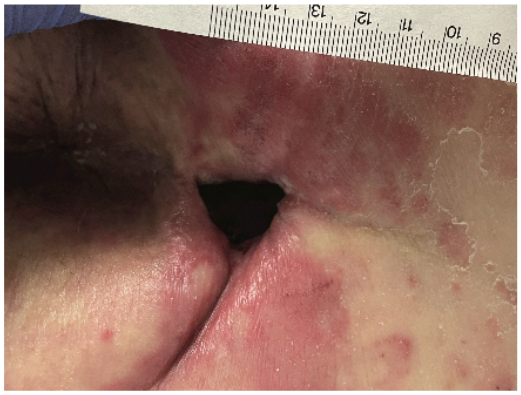

Subject -010

**Supplemental Figure 1: Photos of subject wounds before debridement procedure.**
