## Supplementary material for "What is Slough? A pilot study to define the proteomic and microbial composition of wound slough and its implications for wound healing": Sup. Fig. 2

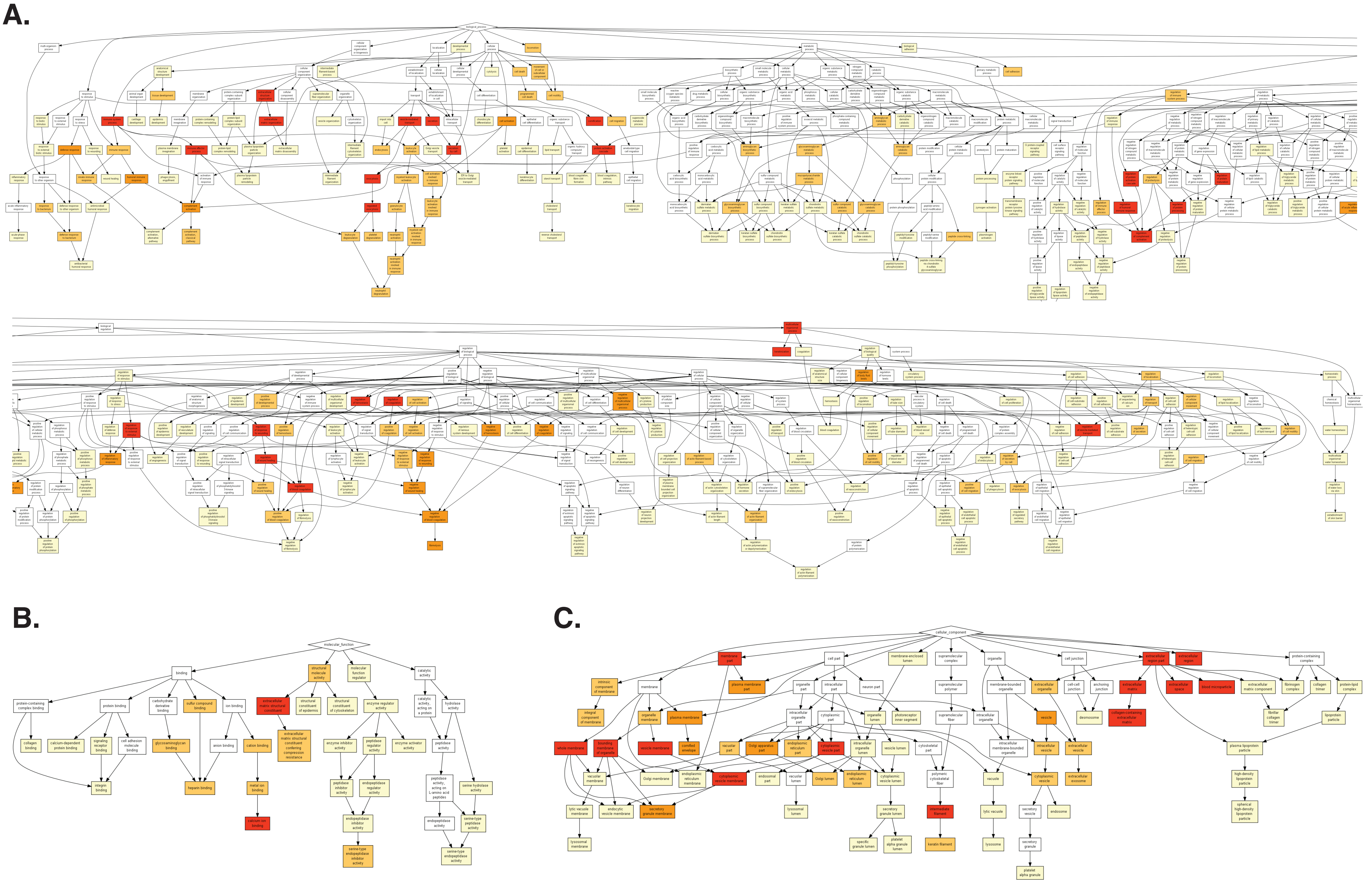


**Supplemental Figure 2: Directed Acyclic Graph (DAG) of the significantly enriched gene ontology (GO) terms grouped by biologic processes (A) molecular functions (B) and cellular components (C) within wound slough.** The most abundant proteins across all slough debridement tissue samples were input as a ranked list to the Gene Ontology enRIchment analysis (GORILA) and visualization tool.27 Figure 1 displays the enriched GO terms associated with each DAG. The significantly enriched GO terms for each DAG are displayed. Box colors indicate p-values; white > 10-3 ; yellow 10-3 – 10-5, yellow-orange 10-5 – 10-7, orange 10-7 – 10-9, Red < 10-9. More detail, including GO term annotations, descriptions, enrichment, number of proteins (Uniprot Genes) involved from our dataset involved in each GO Term, and FDR-qValues are in Supplemental Table 3.
