## Supplementary material for "What is Slough? A pilot study to define the proteomic and microbial composition of wound slough and its implications for wound healing": Sup. Fig. 3

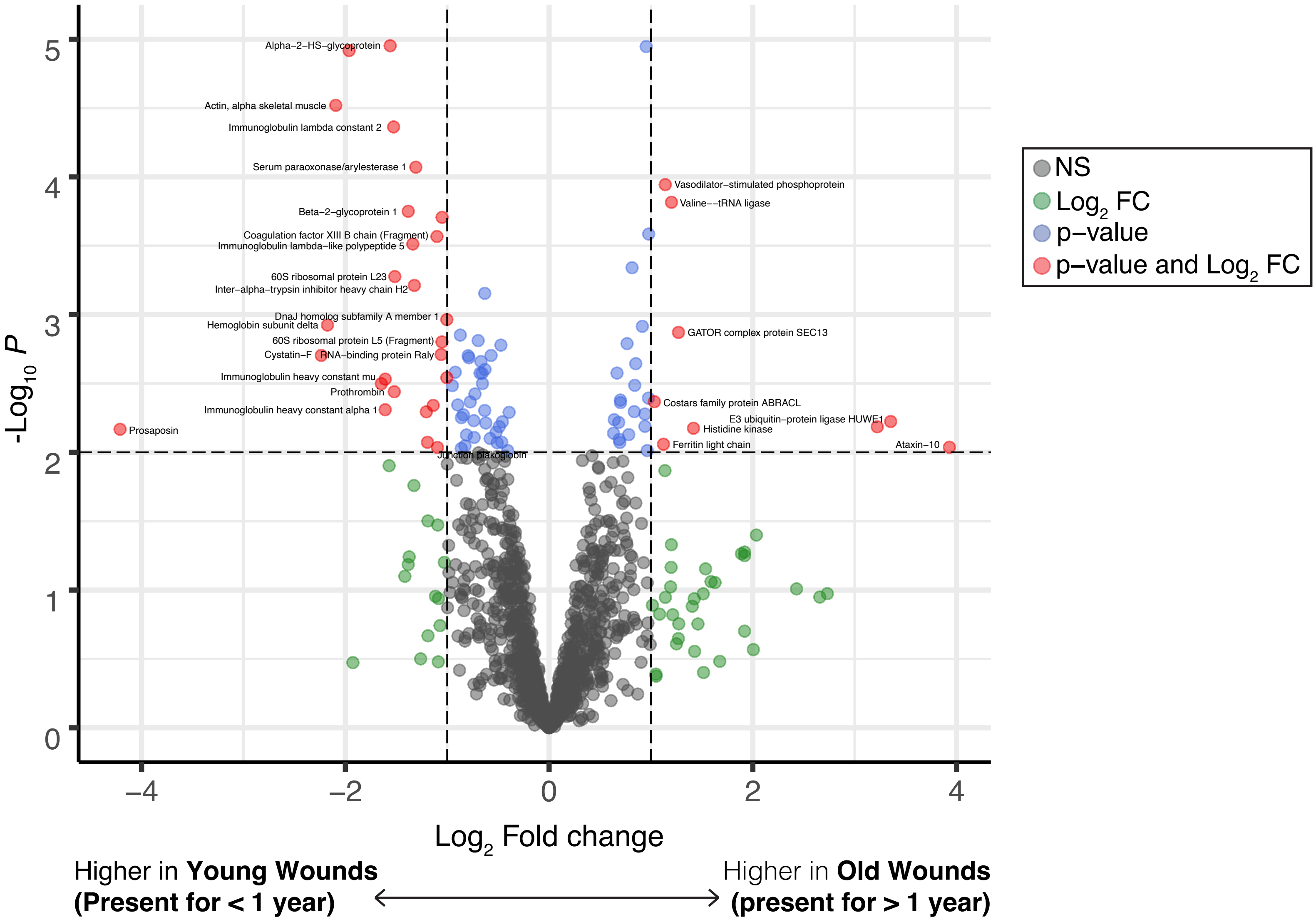


**Supplemental Figure 3: Chronic wounds present less than 1 year are enriched for proteins involved in epithelial barrier formation, neutrophil degranulation, and response to bacteria.** Conversely, wounds present for more than 1 year are enriched for proteins involved in iron sequestration and tRNA metabolism. Subjects were grouped the age of the wound at the time of sample collection. Wounds present for less than 1 year were considered "young", and those present for more than 1 year were considered "old." Groups were assessed for differential protein expression via DEqMS. This volcano plot displays the proteins with significantly greater expression in younger or older wounds.
