## Supplementary material for "What is Slough? A pilot study to define the proteomic and microbial composition of wound slough and its implications for wound healing": Sup. Fig. 4

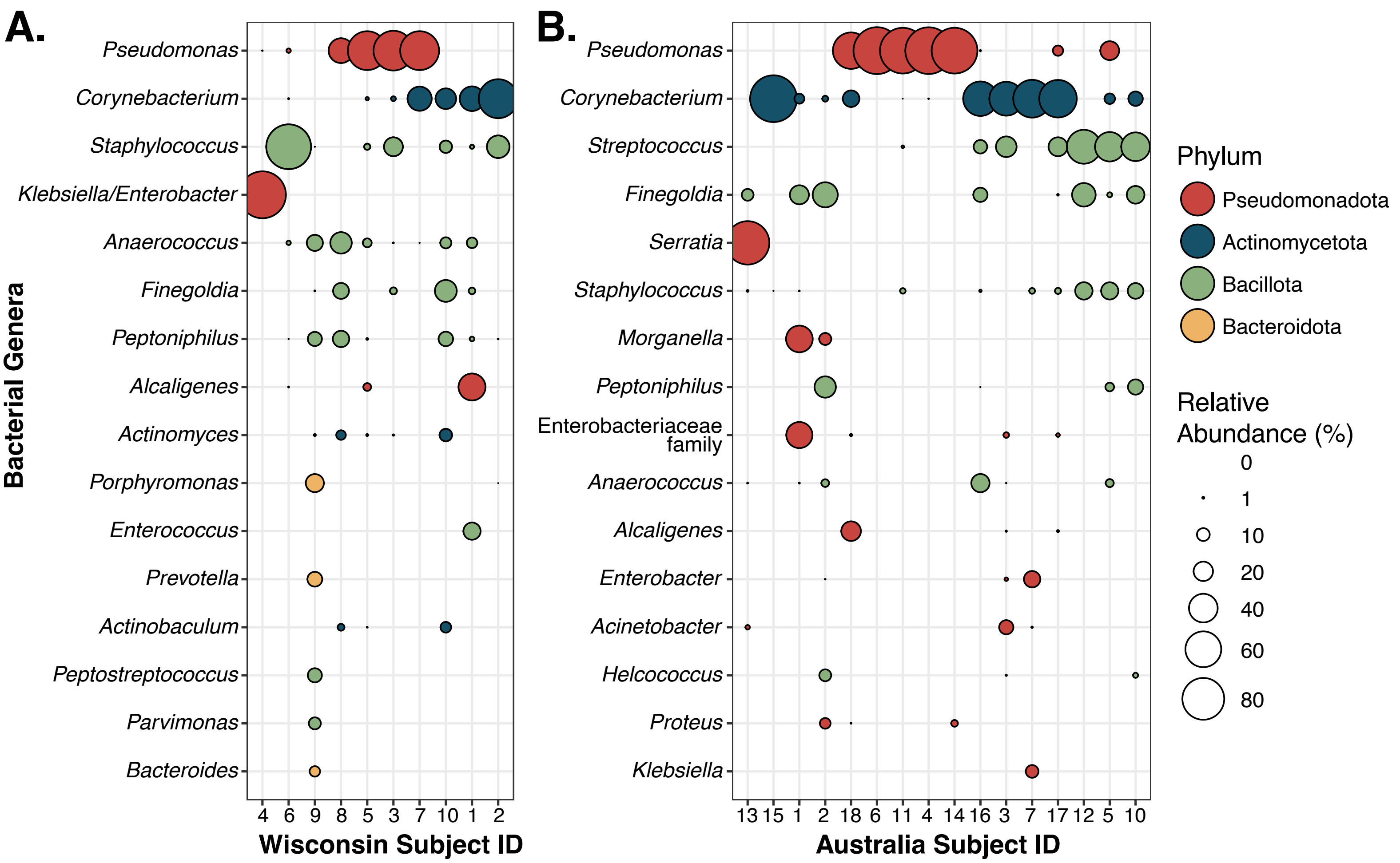


**Supplemental Figure 4: The Abundance of key bacterial taxa is similar across wound slough from two distinct subject cohorts from The United States and Australia.** Datasets were generated using amplicon sequencing of the V4 (panel A, Wisconsin) or V1V3 (panel B, Australia) regions of the 16S rRNA gene, and were thus analyzed separately. ASVs were summed at the genus level. Note that the taxonomic resolution for classification may differ by amplicon region. Genera are shown if present at above 5% relative abundance in at least one specimen and are ordered by mean relative abundance across all specimens within a dataset. Subjects are ordered by average linkage hierarchical clustering of Bray-Curtis dissimilarities. In the Wisconsin cohort (panel A), subject taxa profiles are averaged from multiple specimens.
