## Supplementary material for "What is Slough? A pilot study to define the proteomic and microbial composition of wound slough and its implications for wound healing": Sup. Fig. 5

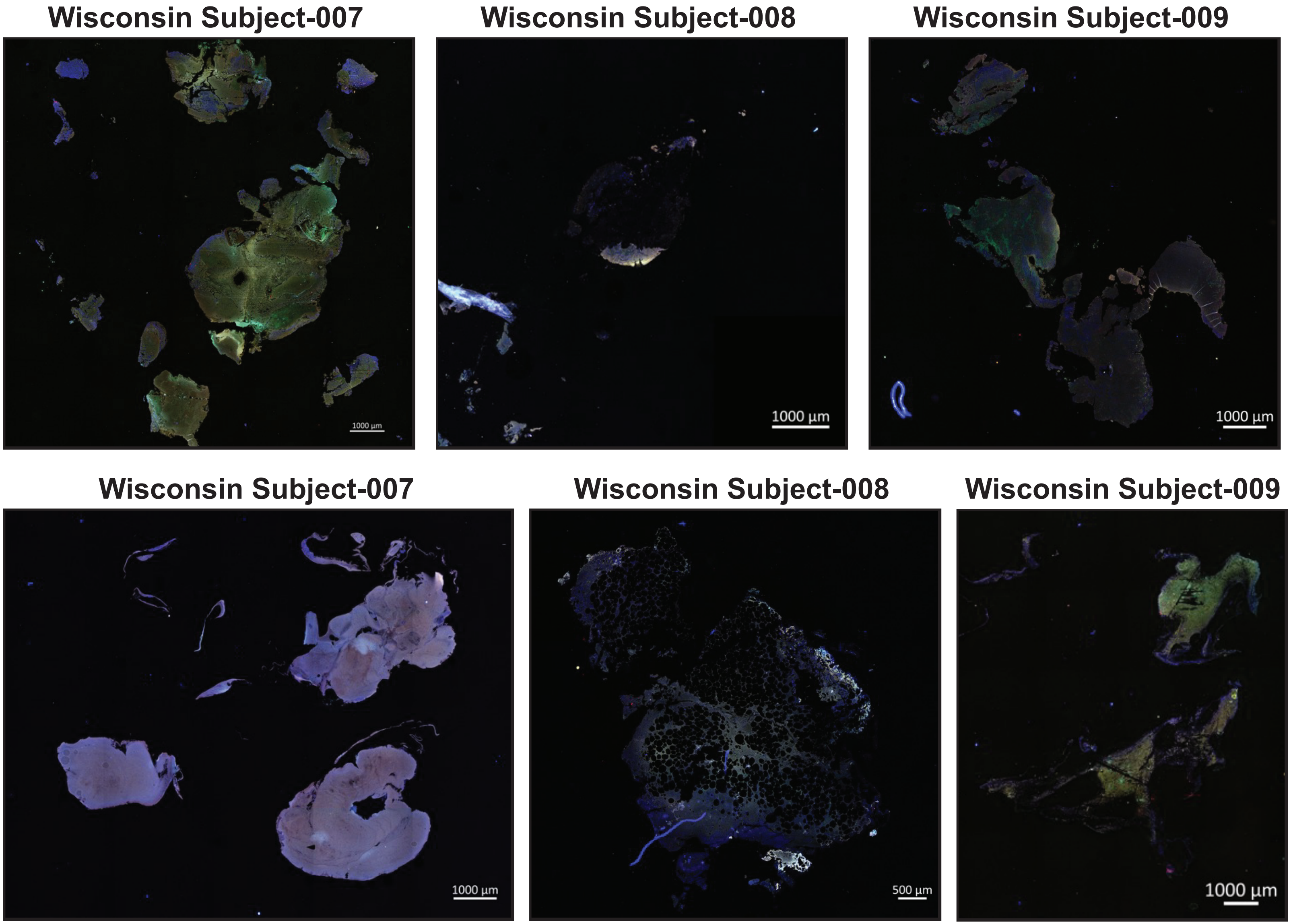


**Supplemental Figure 5: Confocal scanning laser microscopy of slough samples without bacterial aggregates.** Formalin-fixed, paraffin-embedded (FFPE) slough samples were stained with a universal bacterial 16S rRNA probe (red) and for double stranded DNA (DAPI, blue) then visualized with confocal scanning laser microscopy (CSLM). Autofluorescence of the surrounding tissue was visualized in green. The specimens with detected bacterial aggregates are shown in figure 4. This fugure shows the remaining specimens from both patient cohorts that did not have identifiable bacterial aggregates.
