## Supplementary material for "What is Slough? A pilot study to define the proteomic and microbial composition of wound slough and its implications for wound healing": Sup. Fig. 6

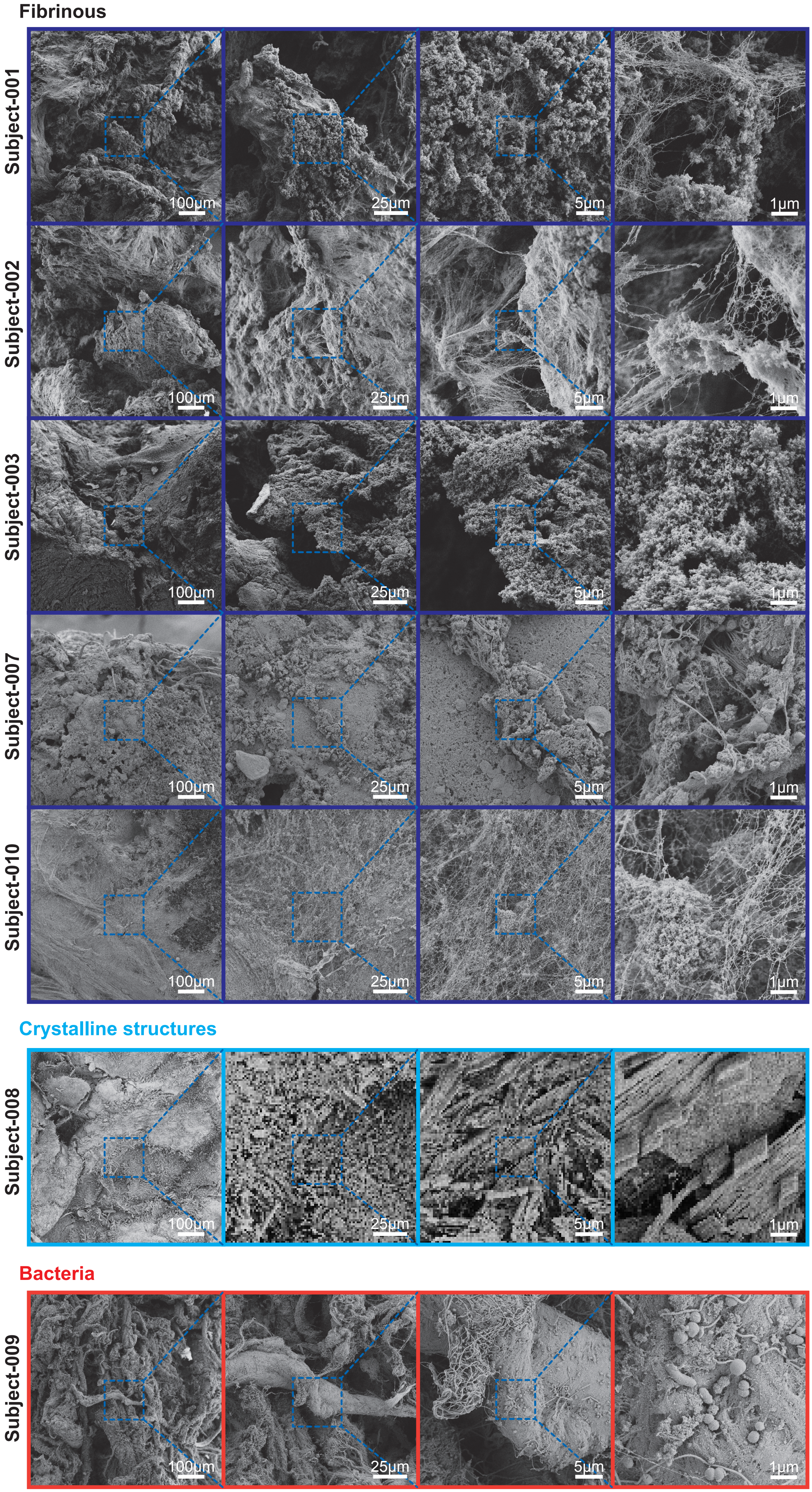


**Supplemental Figure 6: Scanning electron microscopy finds slough to be variable in structure and unique to the subject.** Debrided slough samples were evaluated via scanning electron microscopy (SEM). Subjects-004, -005, and -006 did not have enough debridement tissue for SEM. One subject, subject-009 had visible microorganisms on SEM. A majority of specimens were fibrous in appearance, while one specimen had crystalline structures.
