## Supplementary material for "What is Slough? A pilot study to define the proteomic and microbial composition of wound slough and its implications for wound healing": Sup. Fig. 7

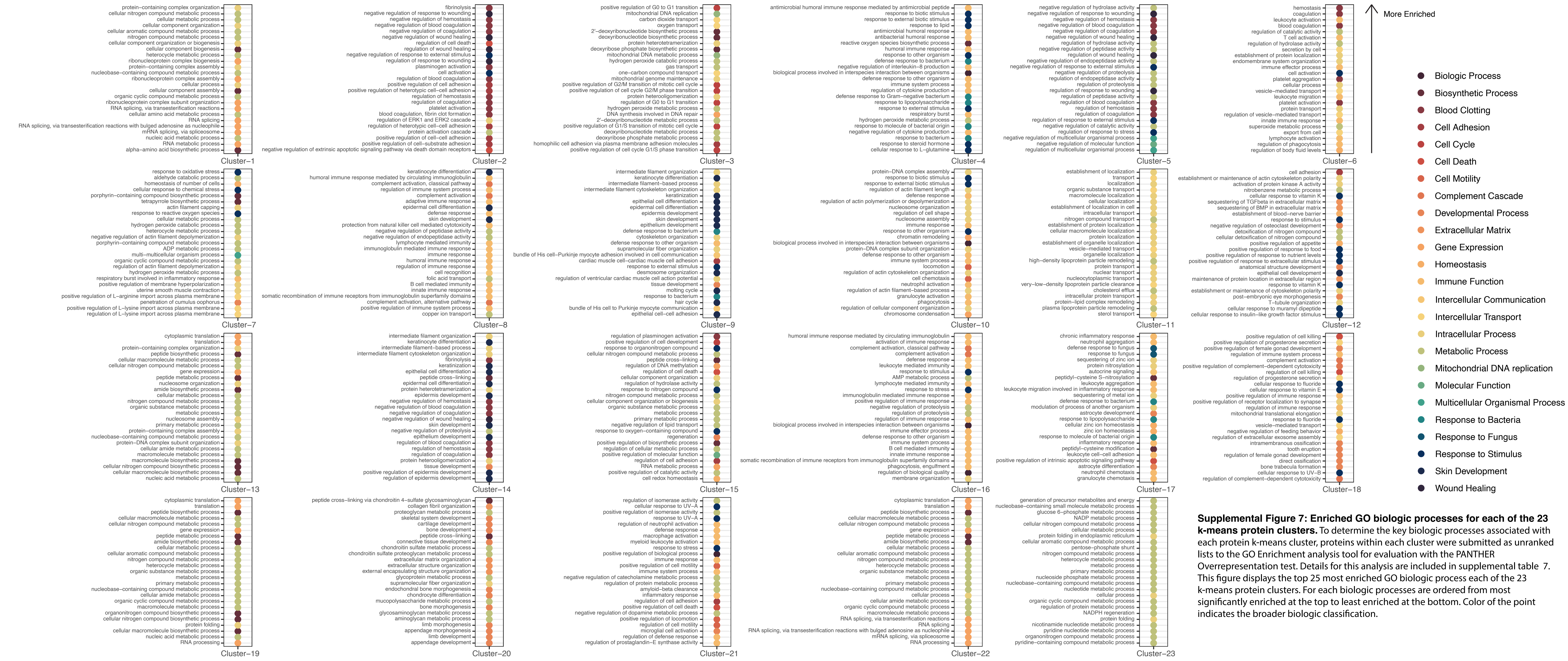


**Supplemental Figure 7: Enriched GO biologic processes for each of the 23 k-means protein clusters.** To determine the key biologic processes associated with each protein k-means cluster, proteins within each cluster were submitted as unranked lists to the GO Enrichment analysis tool for evaluation with the PANTHER Overrepresentation test. Details for this analysis are included in supplemental table 7. This figure displays the top 25 most enriched GO biologic process each of the 23 k-means protein clusters. For each biologic processes are ordered from most significantly enriched at the top to least enriched at the bottom. Color of the point indicates the broader biologic classification.
